# Engineering hyaluronic acid-binding cytokines for enhanced tumor retention and safety

**DOI:** 10.64898/2026.04.06.716711

**Authors:** Elizabeth Fink, William Pinney, Lauren Duhamel, Riyam Al Msari, David Krum, Jordan Stinson, K. Dane Wittrup

## Abstract

Intratumoral delivery of immunotherapy offers a means to enhance efficacy while limiting systemic toxicity, yet rapid diffusion from the tumor constrains dosing levels. Extracellular matrix-targeted anchoring strategies have emerged to improve tumor retention, but the influence of matrix target choice remains poorly understood. Here, we engineered a hyaluronic acid-anchoring platform and directly compared it to a well-established collagen-binding strategy for the delivery of IL-12/IL-15 combination therapy, assessing pharmacokinetic, efficacy, and toxicity endpoints. Hyaluronic acid anchoring markedly enhanced intratumoral retention and tumor loading relative to both unanchored and collagen-anchored constructs. While all anchored cytokine therapies achieved comparable curative tumor control, hyaluronic acid anchoring was associated with improved tolerability, including attenuated systemic inflammation, reduced liver toxicity, and diminished local tissue damage. Analysis of intratumoral immune signaling further indicated that the anchoring strategy modulates local cytokine exposure and immune cell infiltration, despite similar therapeutic outcomes. These findings demonstrate that extracellular matrix target selection significantly shapes the pharmacologic and safety profiles of intratumoral biologics, and identify hyaluronic acid anchoring as an alternative retention strategy with potential advantages.

---

Many immunostimulatory therapies are constrained by narrow therapeutic windows, with doses required for efficacy approaching those that induce systemic toxicity (*1–3*). As a result, considerable effort has focused on restricting drug activity to the tumor, where immune activation or cytotoxicity is desired, while limiting exposure to healthy tissues (*4,5*). Intratumoral administration was proposed to provide such spatial control, but unmodified therapeutic agents readily diffuse out of the tumor and enter systemic circulation, limiting the magnitude of local response while still potentiating on-target, off-tumor toxicity (*6*). To overcome this, a variety of intratumoral anchoring strategies have been developed to retain therapeutic payloads within the tumor microenvironment. Among these approaches, the use of binders targeting extracellular matrix (ECM) components has been widely adopted for use with multiple immunotherapies (*7–11*).

Clinical validation of ECM-targeted anchoring strategies has lagged behind their preclinical success. Most studies to date have targeted a narrow set of ECM components, such as collagen and fibronectin, leaving much of the design space unexplored. While molecular weight and binder affinity predictably determine retention within these systems, it remains unclear how other ECM-specific factors influence the distribution and function of anchored cargo in the tumor microenvironment (*12–14*). This distinction is particularly relevant in the context of tumor heterogeneity, as ECM composition, architecture, and spatial relationships to tumor and immune cells vary both inter- and intratumorally (*15,16*). Thus, the choice of ECM target is likely to influence not only overall retention but also the bioavailability of the therapeutic to target cells.

To explore these open questions, we focused on hyaluronic acid (HA), a highly abundant and metabolically dynamic glycosaminoglycan that is frequently enriched in solid tumors (*17,18*). Compared with collagen, HA occupies a distinct spatial and functional niche within tumors, providing the opportunity to test whether matrix target choice can meaningfully influence therapeutic performance (*19–21*). While collagen-binding strategies have been extensively characterized, HA targeting has been minimally explored, with prior work limited to peptide-binding approaches in isolated therapeutic contexts (*8,22–24*). Here, we engineered an HA-anchoring platform, using targeted protein engineering to optimize binding, stability, and fusion compatibility. We applied this platform to IL-12/IL-15 combination therapy, a regimen with a particularly narrow therapeutic window (*25*). We then compared our HA-anchored cytokines to a well-established, size-matched collagen-anchored system to examine how ECM target influences therapy retention and function. HA anchoring significantly improved tumor retention and loading, which were concomitant with improved tolerability and downstream immune activation. This work establishes HA anchoring as a viable approach for the intratumoral retainment of anticancer therapeutics and underscores the importance of tailoring anchoring strategies to payload properties and tumor architecture to achieve the desired localization, safety, and efficacy.

## RESULTS

### Engineering a high-affinity, stable HA

Several naturally occurring hyaluronic acid binding proteins (HABPs) have been reported and characterized in the literature (*26–28*). To identify one suitable for our anchoring system, we used yeast surface display to screen a panel of HABPs for both stable surface expression and measurable hyaluronic acid binding. This screen yielded a strong candidate, the G1 domain of versican (fig. S1, A to D). Versican is an endogenous proteoglycan that stabilizes extracellular and pericellular HA matrices. While full-length versican has diverse biological functions and binding partners, its G1 domain is specialized for HA binding through its two tandem Link modules (Fig. 1A) (*29,30*).

**Fig. 1.**
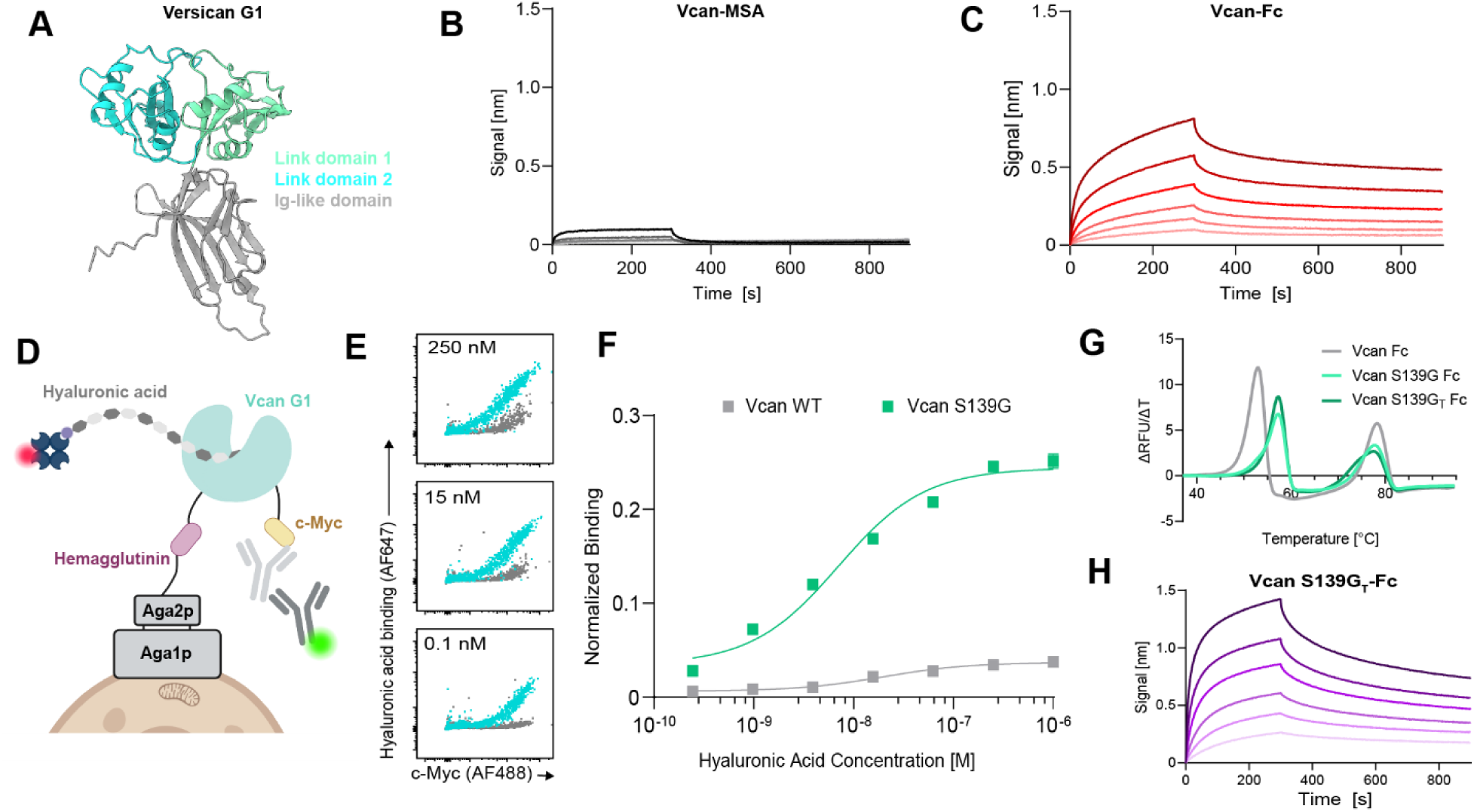
Engineering a stable and high affinity hyaluronic acid anchor. **A**, Alphafold 3 predicted structure of versican G1 domain highlighting the Ig-like domain (gray) and two Link modules (cyan) (AF-Q4VA91-F1-v6). **B, C**, Biolayer interferometry of (**B**) versican G1-MSA and (**C**) versican G1-mIgG2c Fc (LALA-PG) demonstrating association to and dissociation from immobilized hyaluronic acid. **D**, Schematic of yeast surface display system used for affinity maturation of versican G1. Display was detected using a AF488 labeled anti-c-Myc antibody and binding to biotinylated hyaluronic acid was detected using AF647 labeled streptavidin. **E**, Select flow cytometry plots of WT and mutant versican G1 display and binding. Yeast were induced at 37°C and incubated with hyaluronic acid to equilibrium. **F**, Equilibrium binding curves of WT and mutant versican G1 displayed on yeast surface, as in (**E**). Normalized binding was calculated by dividing the binding signal (AF647 MFI) by the display signal (AF488 MFI). **G**, Melting curves of mutant versican-Fc fusion proteins. **H**, Biolayer interferometry of versican S139G_T_-Fc.

We first expressed versican G1 either as a monomeric fusion to mouse serum albumin (MSA) or as a dimeric fusion to an effector-attenuated mouse IgG2c Fc domain (LALA-PG) (fig. S2A) (*31*). Biolayer interferometry revealed that monomeric versican-MSA binds weakly to hyaluronic acid, with a strikingly low signal magnitude and correspondingly poor apparent affinity (K_D_>1 mM) (Fig. 1B). Dimerization of the binding domain increased binding response and yielded an apparent affinity on the order of 100 nM (Fig. 1C). This improvement confirmed that avidity contributes strongly to HA binding, as expected for a highly repetitious polymer. In order to further enhance the intrinsic affinity of the versican HA binding domain we generated a mutagenized library of versican G1 and screened variants with yeast surface display (Fig. 1D). This screen revealed a single mutation, S139G, within the first Link module that substantially increased HA-binding signal (Fig. 1, E and F). A multiple sequence alignment of Link domains across various identified HABPs revealed this to be a reversion-to-consensus mutation (fig. S2B). This, together with higher surface expression for versican S139G relative to wildtype at high temperatures, suggests improved affinity may be conferred by stabilization of the binding domain (fig. S2C) (*32*).

Wildtype and S139 variants were expressed recombinantly as monomeric MSA and dimeric Fc fusions (fig. S2A). The S139G mutation yielded a several log-fold improvement in affinity in both monomeric and dimeric formats (fig. S2, D and E, and Table S1). Melting curve analysis of Fc-fusions confirmed that versican S139G-Fc exhibits greater thermal stability relative to wildtype (Fig. 1G). Subsequently, during construct optimization, we observed poor expression of N-terminal wildtype and S139G fusions. Examination of the structure revealed a short, seven-amino acid disordered segment (L1-E7) of the Ig-like domain at the N terminus that when truncated, rescued expression of N-terminal versican fusions while maintaining improvements in thermal stability and affinity over wildtype versican G1 (Fig. 1, G and H, fig. S2, F and G, and Table S1). The final optimized construct, versican S139G_T_-Fc, is a stable and specific HA-binding anchor suitable for fusion to therapeutic payloads.

### HA anchoring enhances intratumoral retention and tumor loading

We sought to determine whether HA anchoring improves intratumoral persistence and distribution relative to collagen-binding. We evaluated differences in protein pharmacokinetics and biodistribution by comparing our engineered HA-binding protein (versican-S139G_T_-Fc) to a validated collagen-binding Fc fusion (lumican-Fc) and an irrelevant fluorescein-targeted antibody control (fig. S3, A and B) (10). Lumican binds specifically to collagen I and collagen IV, which are frequently overexpressed in the ECM of solid tumors (*10,33,34*).

To quantify intratumoral retention, tumor-localized fluorescence was monitored over time following intratumoral administration of AF647-labeled protein into albino mice bearing B16F10 TRP2-KO tumors (Fig. 2A). The fitted half-lives were 7 hours for the untargeted control antibody, 22 hours for lumican-Fc (>3-fold increase relative to unanchored control), and 73 hours for versican S139G_T_-Fc (>10-fold increase relative to unanchored control) (Fig. 2, B and C). These data indicate that HA anchoring substantially enhances intratumoral persistence compared to both unanchored and collagen-anchored proteins.

**Fig. 2.**
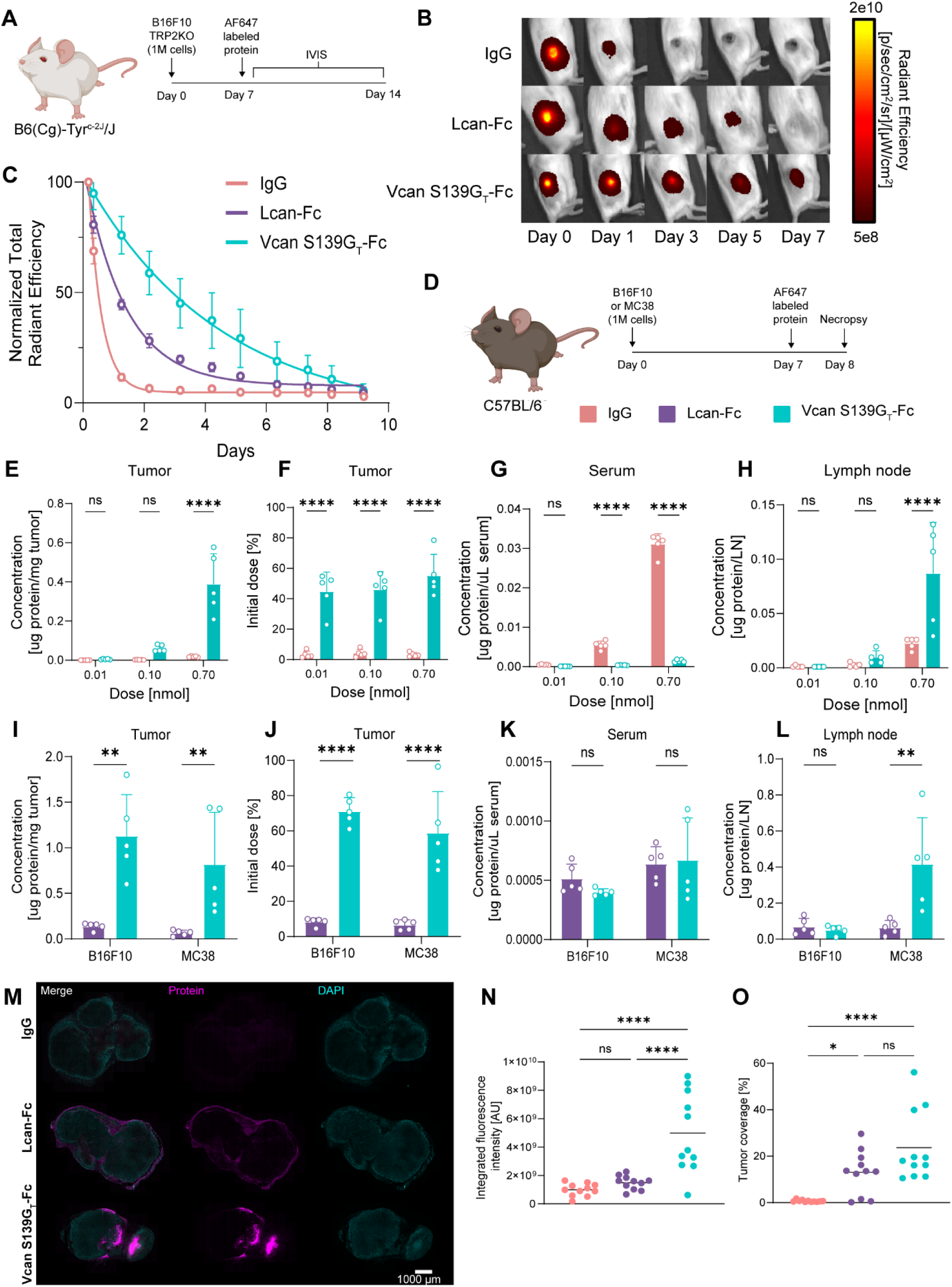
HA anchored protein exhibits a distinct intratumoral spatiotemporal profile. **A**, Albino mice bearing B16F10 TRP2-KO were intratumorally injected with 0.1 nmol of AF647-labeled untargeted IgG, lumican-Fc, or versican S139G_T_-Fc 6 days after tumor inoculation (mean ± SD, n = 4 or 5). Fluorescence was longitudinally tracked with IVIS. **B**, Images and (**C**) quantification of intratumoral retention of constructs. **D**, Mice bearing B16F10 or MC38 flank tumors.were intratumorally injected with indicated dose of fluorescently labeled constructs 6 days after tumor induction. Tissues were harvested for compartmental biodistribution or immunofluorescent microscopy 24 hours later for analysis. **E-H**, Dose escalation experiment was performed in B16F10 tumor bearing mice (mean ± SD, n = 5). **E**, Concentration of IgG and HA-anchored protein in the tumor. **F**, Percent initial dose inside the tumor obtained from protein concentration, tumor weight, and initial dose. **G, H**, Concentrations of IgG and HA-anchored protein in (**G**) serum and (**H**) tdLN. **I-L**, Comparison of HA- and collagen-anchored capacity was performed in MC38 and B16F10 models (mean ± SD, n = 5). **I**, Concentration of collagen- and HA-anchored protein in the tumor. **J**, Percent initial dose of proteins inside the tumor. **K, L**, Protein concentrations in (**K**) serum and (**L**) tdLN. **M-O**, B16F10 tumor-bearing mice were injected intratumorally with fluorescently labeled proteins, and tumors excised 24 hours later for immunofluorescent imaging (n=4, 2-3 slices per tumor). **M**, Representative immunofluorescent microscopy images, (**N**) quantification of integrated fluorescent intensity, and (**O**) percent tumor coverage. Statistics: two-way ANOVA (**E-L**) or one-way ANOVA (**N, O**) with Tukey’s multiple comparison test. ns, not significant; *P < 0.05; **P < 0.01; ****P < 0.0001.

We next assessed whether HA anchoring increases the tumor loading capacity of intratumorally delivered proteins. In the B16F10 model, escalating doses of fluorescently labeled versican S139G_T_-Fc or an untargeted control antibody were administered, and protein levels in the tumor, tumor-draining lymph node (tdLN), and serum were quantified 24 hours post-injection (Fig. 2D). Intratumoral concentrations of versican S139G_T_-Fc were consistently higher than those of the untargeted antibody, and the fraction of the injected dose that was retained within the tumor (%ID) remained stable up to 0.7 nmol (∼90 µg) (Fig. 2, E and F). The maintenance of a constant %ID across increasing doses indicates that tumor-associated HA binding sites were not saturated under these conditions. Systemically, HA-anchored protein was detected at markedly lower concentrations in serum than the unanchored control, while tdLN concentrations increased modestly with escalating doses (Fig. 2, G and H). In total, these results indicate that HA anchoring can safely sequester therapeutics within the tumor microenvironment, and that the loading capacity of HA-anchored therapeutics in B16F10 exceeds a staggering 90 µg.

To determine whether this high tumor retention represents a distinct advantage of HA anchoring, we repeated this analysis at the 0.7 nmol dose in both MC38 and B16F10 tumor models, and compared this treatment to collagen-anchored lumican-Fc. HA-anchored protein remained at significantly higher intratumoral levels after 24 hours (69% of the injected dose in B16F10, 64% in MC38), whereas collagen-anchored lumican-Fc was mostly cleared (5-7% of the injected dose in both tumor models) (Fig. 2, I and J). As expected, little protein was detected outside the tumor for either construct (Fig. 2, K and L). These differences significantly exceed what can be explained by increased retention time alone, highlighting tumor loading capacity as a key factor distinguishing HA anchoring from collagen anchoring in these models. Taken together with our IVIS measurements, these findings demonstrate that HA anchoring has the potential to substantially increase intratumoral cumulative drug exposure (area under the concentration-time curve; AUC) relative to unanchored and collagen-anchored counterparts.

Finally, we examined how HA- and collagen-anchored proteins distribute within the tumor. B16F10 tumor-bearing mice were injected intratumorally with a nonsaturating dose of fluorescently labeled proteins, and their tumors excised 24 hours later for immunofluorescent imaging. Protein fluorescence and DAPI staining were used to quantify integrated fluorescent signal and percent tumor coverage (Fig. 2M and fig. S4, A and B). Versican S139G_T_-Fc exhibited significantly higher total fluorescence within tumors compared with lumican-Fc and the irrelevant control antibody, which aligns with the intratumoral retention data and reinforces that HA anchoring promotes greater tumor-localized protein confinement (Fig. 2N). Both anchored proteins covered a greater fraction of tumor area than the control antibody, although coverage varied across sections, reflecting the spatial heterogeneity of the tumor ECM (Fig. 2O). Despite similar coverages, the anchors imparted qualitatively distinct localization patterns; collagen-anchored protein was distributed more diffusely along tumor borders, whereas HA-anchored proteins formed concentrated depots both within and around the tumor mass (Fig. 2M). Together, these findings highlight how HA-anchoring imparts a distinct spatiotemporal profile relative to another established protein-based tumor localization strategy.

### HA and collagen anchoring similarly enhance IL-12/IL-15 combination efficacy

The observed differences in the biodistribution and pharmacokinetics of these anchoring technologies prompted us to compare them in a therapeutic context. To probe the functional consequences of HA versus collagen anchoring, we selected IL-12 and IL-15 combination therapy, an intratumoral regimen that is highly efficacious yet potently toxic when delivered unanchored and that has been previously characterized by our lab (*35*). We designed a panel of constructs to test the effects of molecular weight and avidity on treatment efficacy and toxicity. This panel included untargeted control constructs (IgG-IL12, IgG-IL15, IL12-VHH_2_-MSA, and VHH_2_-MSA-IL15), HA-binding constructs (IL12-versican S139G_T_-Fc, versican S139G_T_-Fc-IL15, IL12-versican S139G_T_-MSA, and versican S139G_T_-MSA-IL15), and collagen-binding constructs (IL12-lumican-Fc, lumican-Fc-IL15, IL12-lumican-MSA, and lumican-MSA-IL15) (Fig. 3A and fig. S5, A and B). All constructs retained binding towards their respective ligands and preserved cytokine activity (fig. S5, C to I).

**Fig. 3.**
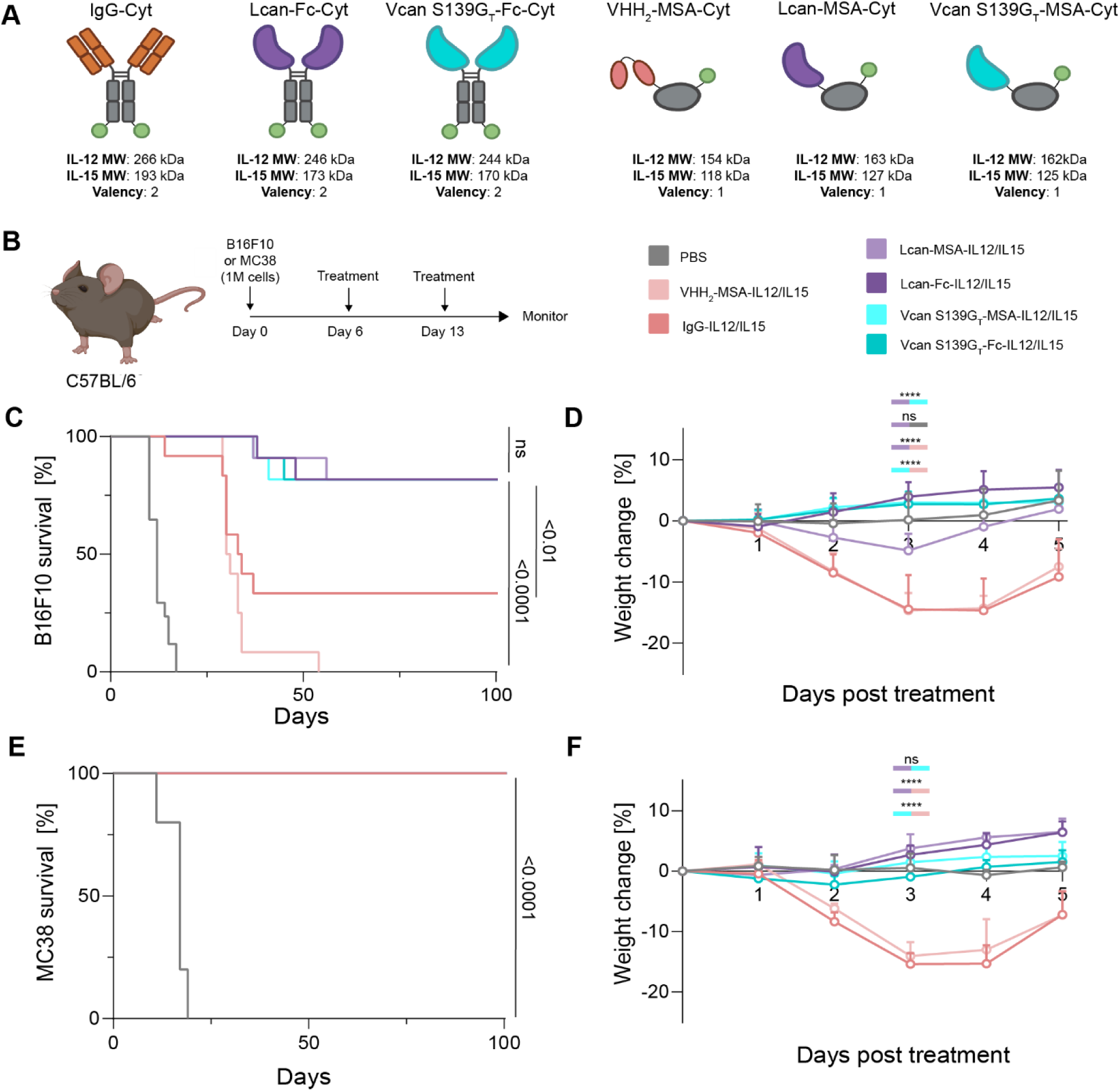
Collagen- and HA-anchored IL-12/IL-15 combination therapies are equally efficacious. **A**, A panel of dimeric (Fc) and monomeric (MSA) cytokine fusions to untargeted, collagen-anchored, or HA-anchored proteins was engineered. **B**, Mice bearing B16F10 or MC38 tumors were treated days 6 and day 13 after tumor induction with 0.11 nmol IL-15 and 0.014 nmol IL-12. Mice were monitored for tumor burden and weight loss. **C**, Kaplan-Meier survival and (**D**) weight loss of mice bearing B16F10 tumors intratumorally injected with PBS or cytokine fusion proteins (mean ± SD, n = 11). **E**, Kaplan-Meier survival and (**F**) weight loss of mice bearing MC38 tumors treated with PBS or cytokine fusions (mean ± SD, n = 5). Statistics: analysis of survival was performed using log-rank Mantel-Cox test. Weights compared by two-way ANOVA with Tukey’s multiple comparison test. ns, not significant; ****P < 0.0001.

Mice bearing established B16F10 tumors received intratumoral doses of dual cytokine treatment on days 6 and 13 post-tumor induction (Fig. 3B). As expected, unanchored cytokines drove substantial toxicity and resulted in suboptimal therapeutic efficacy (Fig. 3, C and D). By contrast, all anchored cytokine formats elicited robust antitumor responses, with 80% of mice achieving complete tumor clearance (Fig. 3C and fig. S6, A to F). Although prior experiments revealed that HA anchoring dramatically improves intratumoral protein retention relative to collagen anchoring, collagen-anchored constructs as well as smaller, monomeric fusions, achieved comparable tumor regression. This indicates that extended intratumoral persistence, while achievable with HA anchoring, is not necessary for therapeutic effect in this context. Despite similar antitumor efficacy, anchor-dependent differences in acute toxicity were evident with this therapy. Treatment with lumican-MSA-IL12/IL15 combination induced pronounced weight loss 3 days after treatment, whereas no weight loss was observed in mice treated with the corresponding HA-anchored monomeric cytokines (Fig. 3D). This toxicity is consistent with systemic leakage of collagen-anchored cytokines, which is minimized in HA-anchored constructs.

To assess whether these effects were tumor-model specific, we repeated the study in the more immunogenic MC38 tumor model using the same dosing and treatment schedule. All treated mice achieved complete tumor regression, consistent with MC38’s higher baseline immune infiltration and greater responsiveness to therapy compared to B16F10 (Fig. 3E and fig. S6, G to L) (*36*). Mice receiving unanchored cytokine therapy experienced significant weight loss, and no significant differences in weight loss were observed across anchored treatment groups (Fig. 3F). The absence of weight loss in lumican-MSA-IL12/IL15 treated mice may reflect ECM differences in MC38 tumors that improve retention and confinement of collagen-anchored proteins relative to B16F10 (*34,37*).

### HA anchoring improves tolerability at high doses

The toxicity seen with collagen-anchored IL-12/IL-15 therapy in the B16F10 model prompted us to further investigate potential anchor-dependent differences in toxicity and to test if HA anchoring could broaden the therapeutic window of this potent combination. We hypothesized that doubling the dose of cytokines would exacerbate the therapy-induced toxicity resulting from lumican-MSA-IL12/IL15 combination. B16F10 tumor-bearing mice were treated once with monomeric anchored or unanchored cytokine fusion 6 days after tumor induction, and body weight was monitored daily as a proxy for systemic toxicity (Fig. 4A). Consistent with our hypothesis, mice receiving unanchored or collagen-anchored cytokine therapy experienced significant weight loss relative to untreated control mice. Notably, versican S139G_T_-MSA-IL12/IL15 combination therapy did not induce weight loss, even at this high dose (Fig. 4, B and C).

**Fig. 4.**
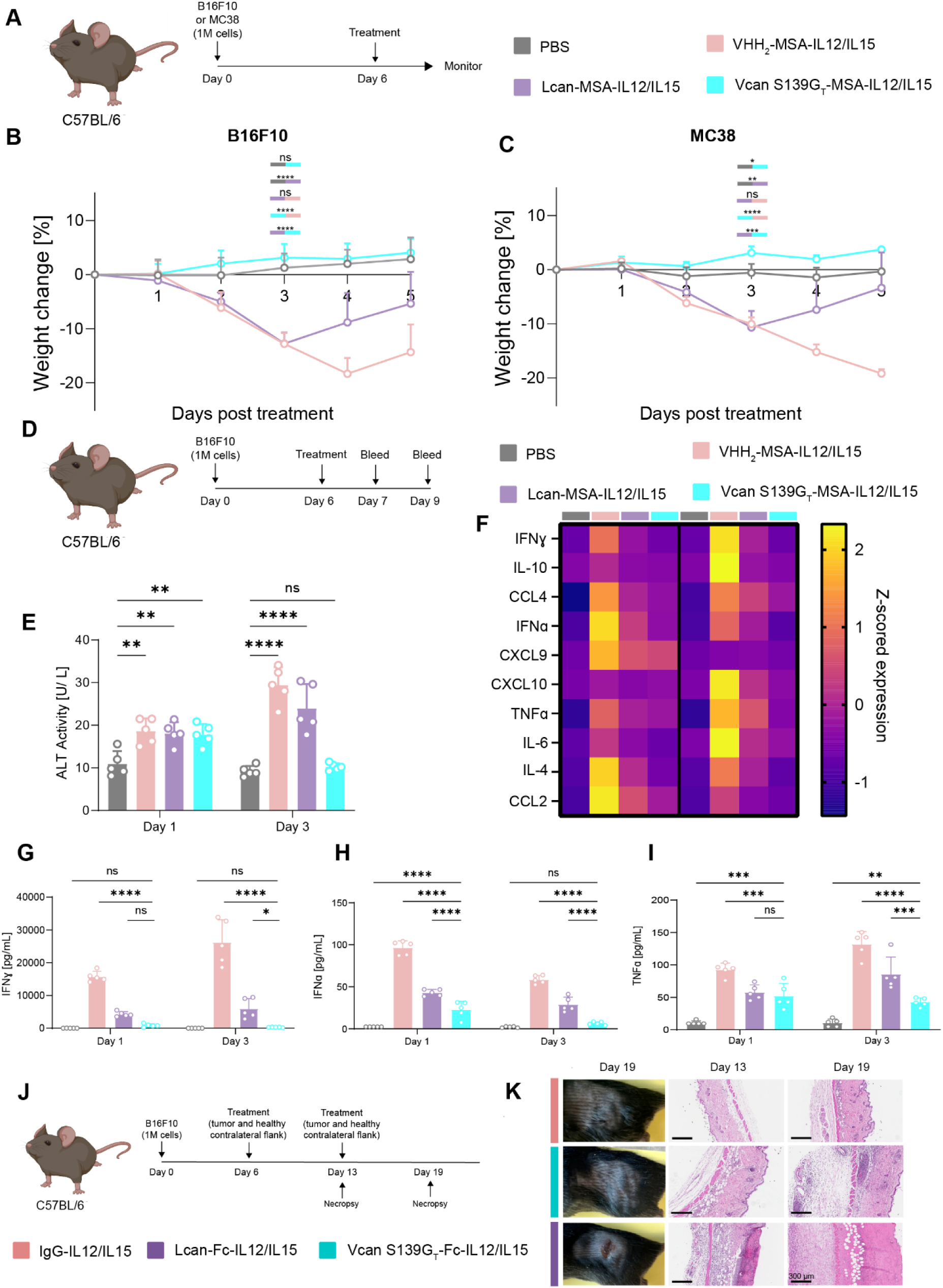
HA-anchored cytokine therapy demonstrates improved tolerability. **A**, B16F10- or MC38-tumor bearing mice were intratumorally injected with 0.22 nmol IL-15 and 0.028 nmol IL-12 fusion proteins 6 days after tumor induction. Weight loss was monitored for 5 days post-treatment (mean ± SD, n =5). Percent weight change measured from the start of treatment in B16F10 (**B**) and MC38 (**C**) tumor-bearing mice. **D**, Mice were treated as described above, and blood collected 1 and 3 days post-treatment for analysis of ALT activity and serum cytokine/chemokine levels (mean ± SD, n =5). **E**, Alanine aminotransferase activity in serum 1 and 3 days post-treatment. **F**, Z-scored expression of serum cytokines and chemokines for each cohort 1 and 3 days post-treatment. **G-I,** Serum concentrations of (**G**) IFNɣ, (**H**) IFNɑ, and (**I**) TNFɑ tested at both timepoints. **J**, B16F10 tumor-bearing mice were treated with 0.11 nmol IL-15 and 0.014 nmol IL-12 of indicated cytokine fusions 6 days post-tumor inoculation. Mice were injected both intratumorally and on their healthy contralateral flank. Seven days after the primary injection, mice were either euthanized for tissue collection or treated again. Six days after the second injection, the rest of the mice were euthanized for histopathological analysis of tissues (n=6). **K**, (Left) Macroscopic images of dosed contralateral flank after 2 cytokine injections. (Center, right) H&E stained skin sections of contralateral flank after 1 (center) or 2 (right) subcutaneous injections of cytokine fusions. Statistics: two-way ANOVA with Tukey’s multiple comparison test. ns, not significant; *P < 0.05; **P < 0.01; ***P < 0.001; ****P < 0.0001.

To further characterize systemic toxicity, we repeated this dosing paradigm and collected serum 1- and 3-days post-treatment (Fig. 4D). Serum alanine aminotransferase (ALT) levels were measured as an indicator of liver toxicity, and circulating cytokines and chemokines were quantified to assess systemic inflammatory responses. One day after treatment, all cytokine-treated groups showed modest elevations in ALT compared to untreated control mice. By day 3, ALT levels in mice receiving HA-anchored cytokines had returned to baseline, whereas mice treated with unanchored or collagen-anchored IL-12/IL-15 exhibited further increases in ALT (Fig. 4E). Consistent with these findings, LegendPlex analysis revealed elevated levels of circulating inflammatory cytokines and chemokines in mice treated with unanchored or collagen-anchored IL-12/IL-15 compared with untreated or HA-anchored groups (Fig. 4F and fig. S7, A to G). Of note, collagen-anchored cytokine therapy elicited increased systemic IFNɣ, IFNɑ, and TNFɑ over HA-anchored cytokine therapy at both timepoints tested (Fig. 4, G to I). Collectively, these data indicate that HA anchoring confers improved systemic tolerability relative to collagen anchoring at high cytokine doses.

In addition to these systemic effects, HA anchoring prevented localized adverse reactions to IL-12/IL-15 therapy. Mice treated with lumican-Fc-cytokine therapy developed skin lesions at the injection site, which were absent in mice receiving unanchored or HA-anchored IL-12/IL-15 therapy (fig. S8, A to C). To further investigate these differences, mice were injected both intratumorally on their tumor-bearing flank and subcutaneously on the healthy contralateral control flank on days 6 and 13 post-tumor induction, and tissues collected on days 13 or 19 (Fig. 4J). Lesions were observed on both the treated tumor and the contralateral flank of lumican-Fc-IL12/IL15 treated mice but not on mice receiving versican S139G_T_-Fc-IL12/IL15. Histological analysis of treated healthy flank revealed that unanchored and HA-anchored cytokines induced only mild to moderate inflammation and edema, whereas collagen-anchored cytokine therapy caused severe inflammation, vascular damage, and eventual local tissue necrosis (Fig. 4K). Although the precise mechanism for this observation remains unclear, our findings indicate that both anchors modulate a local inflammatory response but produce distinct tissue damage profiles (*38*).

### Anchoring shapes the intratumoral immune landscape

Our findings demonstrate that fusing cytokines to half-life extension moieties (Fc and MSA) or different tumor-retention technologies (HA and collagen) alters cytokine pharmacokinetics in ways that can modulate therapy-induced toxicity. Although our anchored cytokine therapies achieved comparable antitumor efficacy, their distinct retention and toxicity profiles raised the question of whether immune cells within the tumor experience differences in local cytokine exposure, and whether such differences are reflected in the intratumoral immune landscape. To address this, we focused on a single cytokine, IL-12, to avoid confounding effects from dual cytokine signaling. Downstream activity was assessed by measuring STAT4 phosphorylation (pSTAT4), a proximal readout of IL-12 receptor engagement, allowing us to link observed immune responses to differences in cytokine biodistribution.

B16F10 tumor-bearing mice were treated intratumorally with PBS or monomeric cytokine fusions, and tumor immune infiltrates were analyzed 5 days post-treatment (Fig. 5A). Among the therapies tested, only HA-anchored IL-12 induced significant increases in both the proportion and count of infiltrating leukocytes relative to PBS, whereas collagen-anchored and unanchored cytokines failed to achieve comparable changes (Fig. 5, B and C). Within the CD45^+^ population, the fraction of pSTAT4^+^ cells was elevated in all treated groups, but trended downward with increasing intratumoral retention, perhaps due to dilution by infiltrating cells not expressing IL-12R (Fig. 5, C to E). Despite this trend, only IL12-versican S139G_T_-MSA produced a significant increase in the absolute number of pSTAT4^+^CD45^+^ cells, an effect not seen with collagen anchoring (Fig. 5F). Similar patterns were observed within the CD3^+^ compartment; where only HA-anchoring yielded significant increases in both CD3^+^ and pSTAT4^+^CD3^+^ tumor infiltrating cells, whereas collagen-anchored IL-12 again failed to reach significance in these metrics (Fig. 5, G to K). As in the CD45^+^ population, the proportion of pSTAT4^+^CD3^+^ T cells decreased with enhanced IL-12 retention (Fig. 5J). Overall, the data suggest that HA anchoring can subtly influence the tumor-infiltrating immune landscape relative to collagen anchoring primarily through increased leukocyte and T cell infiltration, though the potent IL-12/IL-15 combination renders these differences largely inconsequential for the overall therapeutic effect.

**Fig. 5.**
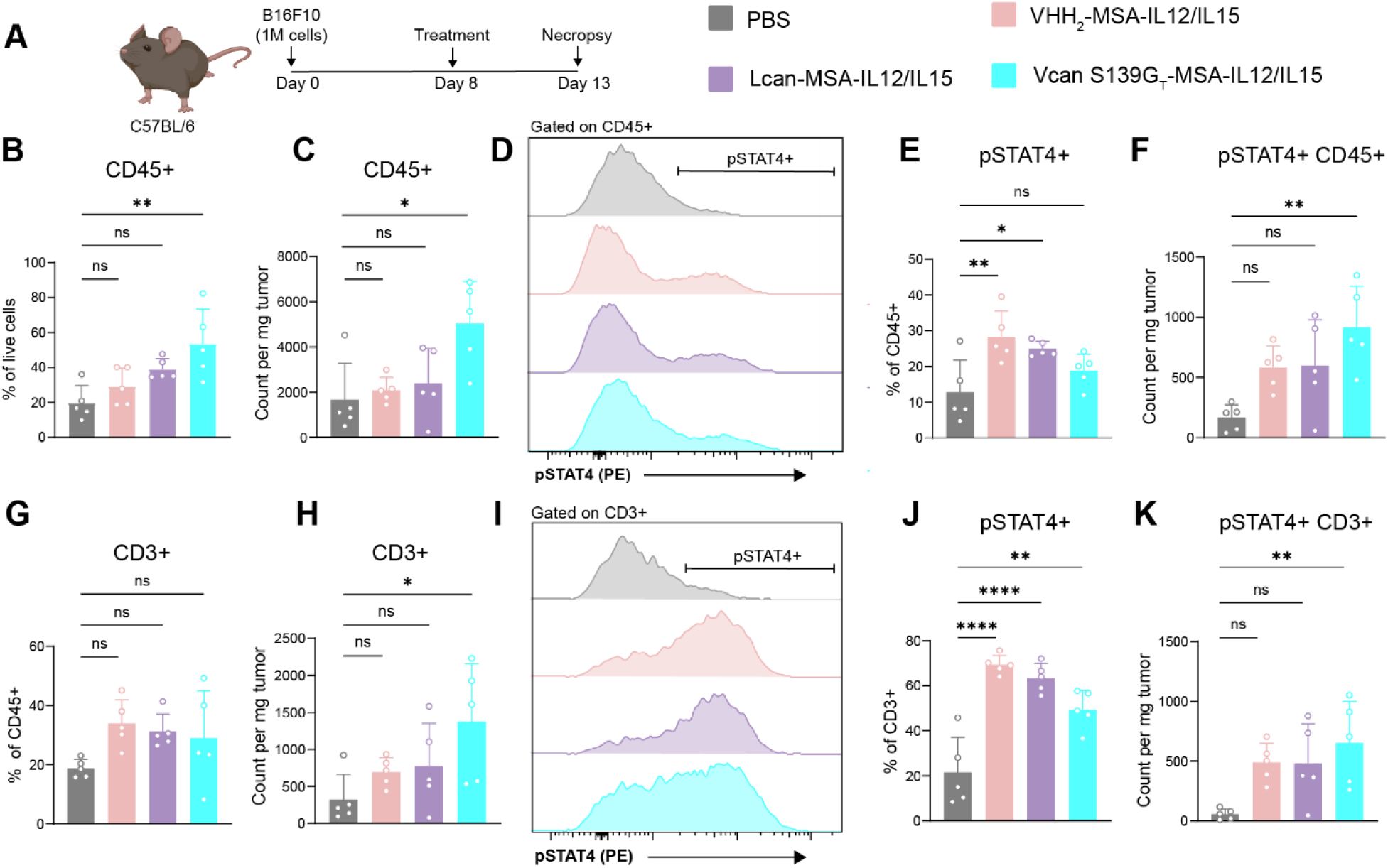
Immune infiltrate and pSTAT4 signaling resultant from anchored IL-12 administration. **A**, B16F10 tumor-bearing mice were intratumorally injected with 0.05 nmol of indicated IL-12 fusion proteins 8 days after tumor inoculation. Mice were euthanized and tumors collected 5 days later for downstream flow cytometric analysis (mean ± SD, n=5). **B,** Treatment effects on proportion and (**C**) count of tumor-infiltrating CD45+ leukocytes. **D**, Representative histograms and gating for pSTAT4+ cells, previously gated on CD45+ cells. **E**, Proportion and (**F**) count of pSTAT4+ cells in the whole CD45+ compartment. **G**, Treatment effects on proportion and (**H**) count of tumor-infiltrating T cells. **I**, Representative histograms and gating of pSTAT4+ cells in the CD3+ compartment. **J**, Proportion and (**K**) count of pSTAT4+ T cells in the tumor. Statistics: one-way ANOVA followed by Tukey’s multiple-comparison test. ns, not significant; *P < 0.05; **P < 0.01; ***P < 0.001; ****P < 0.0001.

## DISCUSSION

In this study, we engineered a stabilized, high-affinity hyaluronic acid-binding protein and used it to evaluate HA anchoring as a strategy for intratumoral cytokine delivery. Although intratumoral administration and ECM-targeted anchoring have each been explored as a means to improve the therapeutic index of immunotherapies, there has been minimal exploration of HA-based anchoring. Here, we establish HA anchoring as a modular, protein-based tumor-localization technology and directly compare it to a well-characterized collagen-binding strategy across pharmacokinetic, therapeutic, and immunologic measurements.

Relative to collagen anchoring, HA anchoring conferred a distinct intratumoral pharmacokinetic profile characterized by prolonged retention (despite HA’s more rapid turnover rate relative to collagen) and significantly increased tumor loading potential (*39*). These findings support that ECM targets are not interchangeable with respect to quantitative tumor retention metrics and that the spatial organization and biochemical properties of the specific matrix components can impart qualitatively different modes of tissue localization. In this context, HA anchoring appears particularly well suited for achieving improved intratumoral drug AUC, likely reflecting the abundance and dynamic distribution of HA within the tumor microenvironment.

Downstream of this enhanced tumor exposure to cytokine, analysis of intratumoral STAT4 phosphorylation in response to IL-12 administration revealed anchoring-dependent differences in cytokine signaling and immune cell infiltration, indicating that anchoring can modulate local cytokine exposure. The improved retention of HA-anchored therapy also had a clear impact on toxicity. HA-anchored cytokines were better tolerated than collagen-anchored counterparts, as evidenced by reduced systemic inflammatory cytokine levels and more rapid normalization of serum ALT. Further, histological assessment of treated healthy tissue revealed collagen-anchored cytokines induced localized vascular damage and tissue necrosis. One possible explanation is that collagen-anchored cytokines may preferentially partition onto collagen IV in the perivascular basement membrane, thereby exacerbating local vascular damage (*40*). These results support that intratumoral retention alone is insufficient to predict safety; rather, the anatomical localization of a therapy among vascular, stromal, and pericellular compartments can substantially influence local toxicity. While our data show that the pharmacokinetic advantages conferred by HA anchoring can meaningfully impact downstream immune response and toxicity, HA anchoring did not improve antitumor efficacy relative to collagen anchoring in the setting of IL-12/IL-15 combination therapy, given the exceptional potency of this cytokine regimen.

These findings raise questions that warrant future exploration. While we demonstrate that HA and collagen anchoring confer distinct pharmacokinetic and toxicity profiles, we do not directly resolve the microscopic spatial distribution of anchored cytokines relative to vascular, stromal, and immune compartments. Higher-resolution spatial approaches, such as multiplexed immunofluorescence, spatial transcriptomics, or intravital imaging, would be informative in exploring how anchor-dependent localization governs cell-specific cytokine exposure and downstream toxicity. Further, our studies focused only on IL-12 and IL-15, which limited the ability to detect therapeutic gains arising from improved intratumoral retention. Evaluation of HA anchoring with other payloads may better reveal contexts in which increased tumor AUC translates into enhanced antitumor activity. Finally, these studies were performed in a limited set of murine syngeneic transplant tumor models, and the extent to which tumor-specific ECM composition or stromal architecture influences anchoring behavior remains to be determined.

Taken together, these results underscore that tumor anchoring is not a binary design choice, but a tunable parameter that influences cytokine pharmacokinetics, immune engagement, and toxicity. HA anchoring represents a distinct and complementary approach to existing matrix-targeting strategies, enabling unusually high tumor loading and prolonged retention without exacerbating systemic or local adverse effects. While these properties did not enhance the efficacy of a highly potent IL-12/IL-15 regimen, they may be advantageous for therapeutic payloads that require higher intratumoral concentrations or extended residence times to achieve maximal activity. More broadly, our findings highlight the importance of matching anchoring strategy to payload properties and therapeutic goals when designing intratumoral biologics.

## MATERIALS AND METHODS

### Study design

This study aimed to develop a hyaluronic acid-anchoring platform for the intratumoral delivery of immunotherapeutic cargo, and to compare the pharmacokinetic behavior, efficacy, and tolerability of hyaluronic acid-anchored cytokines against a well-established, collagen-binding anchoring strategy. Proteins were engineered and expressed in mammalian systems, followed by biochemical and biophysical characterization to confirm binding and activity.

*In vivo* experiments were performed using syngeneic murine tumor models to assess tumor localization, pharmacodynamic signaling, and therapeutic efficacy. Tumor-bearing mice were randomized into treatment groups based on tumor area to minimize baseline bias. Investigators were not blinded during treatment administration or analysis. We did not use a power analysis to calculate sample size. Experimental cohorts consisted of 5 or 6 mice per group, and efficacy and toxicity experiments in the B16F10 model were repeated twice and results pooled for analysis.

### Cell lines

B16F10 (ATCC), MC38 (ATCC), Expi293F (Gibco), HEK-Blue IL-12 (Invivogen), and CTLL-2 (ATCC) cells were cultured following vendor instructions. B16F10-TRP2 KO cells were generated and maintained as previously described (*41*). B16F10, B16F10-TRP2 KO, MC38, and HEK-Blue IL-12 cells were cultured in Dulbecco’s Modified Eagle Medium (DMEM; ATCC) supplemented with 10% fetal bovine serum (FBS; Gibco) and 100 U/mL penicillin-streptomycin (Gibco). CTLL-2 cells were cultured in Roswell Park Memorial Institute 1640 medium (RPMI; ATCC) supplemented with 10% FBS, 100 U/mL penicillin-streptomycin, 10% T-stim with concanavalin A (Corning), 1x sodium pyruvate (Thermo Fisher Scientific), 1x nonessential amino acids (Thermo Fisher Scientific), and 1x beta-mercaptoethanol (Thermo Fisher Scientific). Expi293F cells were cultured in Expi293 Expression Medium (Gibco). All cells were cultured at 37°C and 5% CO_2_. All cells were confirmed to be pathogen-free via mycoplasma and IMPACT testing.

### Mice

Female B6 mice were purchased from Taconic Biosciences (C57BL/6NTac), and female albino B6 mice were purchased from Jackson Laboratory (B6(Cg)-Tyr^c-2J^/J). All mice were 6-8 weeks old at study initiation. All mouse studies were conducted in accordance with local, state, and federal guidelines under the approval of the Massachusetts Institute of Technology Committee on Animal Care.

### Cloning and protein purification

Gene blocks (gBlock; IDT) encoding recombinant proteins were cloned into the gWiz expression vector (Genlantis) using InFusion cloning (Takara Bio) or NEBridge Golden Gate Assembly Kit BsaI-HF v2 (New England BioLabs). Assembled plasmids were transformed into Stellar Competent Cells (Takara Bio), and single clones were isolated. Sequences were verified by full plasmid sequencing (Primordium). Plasmid DNA was isolated from bacterial cultures using NucleoBond Xtra Midi endotoxin-free midi-prep kit (Takara Bio). The untargeted IgG used in this study was an anti-FITC antibody (clone 4-4-20), and the untargeted VHH was a binder to a *Toxoplasma gondii* kinase (clone 1B7). All protein sequences used in this study can be found in Supplementary Table S2.

Transfection of Expi293F cells was performed using protocols adapted from Zhou et. al., 2023 (*42*). Cell cultures were transfected with 1 mg/L plasmid DNA diluted in Opti-MEM (Gibco) using 8 mg/L polyethylenimine (Polysciences) in Opti-MEM. Twenty-four hours post-transfection, cultures were supplemented with 25 mL/L of 20% (w/v) Soy C-CELL S146 Peptone (Organotechnie) in Expi293F medium and 8 mL/L of 500 mM valproic acid (Sigma) in PBS (Corning). Six days post-transfection, cultures were centrifuged at 4,300 x g and supernatants were sterile-filtered, followed by addition of 1/10 volume of 10X PBS (Corning). Fc-fused and MSA-fused (His-tagged) proteins were purified using rProtein A Sepharose Fast Flow resin (Cytiva Life Sciences) and Ni Sepharose High Performance Resin (Cytiva Life Sciences), respectively, according to vendor instructions. Eluted proteins were buffer-exchanged into PBS using Amicon Ultra Centrifugal Filter units (MilliporeSigma) and further purified by size-exclusion chromatography on a Superdex 200 Increase 10/300 GL column (Cytiva Life Sciences) using an AKTA FPLC system (Cytiva Life Sciences). Recombinant proteins were assessed for size and aggregate by sodium dodecylsulfate-polyacrylamide gel electrophoresis and confirmed to contain <0.1 EU per dose using the Endosafe^TM^ Nexgen PTS system (Charles River Laboratories). Purified proteins were aliquoted, flash-frozen in liquid nitrogen, and stored at -80°C until use.

### Yeast surface display and affinity maturation

Yeast surface display, error-prone library generation, and library screening were performed using standard materials and methods as described elsewhere (*43*). For display of wildtype hyaluronic acid binding proteins (HABPs), gBlocks encoding for murine CD44, CD44-TSG6 Link chimera, HAPLN1, and versican G1 were co-transformed with NheI/BamHI (New England Biolabs) digested pCHA backbone into chemically competent RJY100 using the Frozen-EZ Yeast Transformation II Kit (Zymo Research). Transformations were plated on SD-CAA plates and single clones were isolated and sequence-verified. Cultures were grown in SD-CAA at 30°C and induced in SG-CAA overnight at 20°C. All staining of yeast was performed in PBS with 0.1% (w/v) bovine serum albumin (BSA; Sigma Aldrich). Post-induction, yeast cell surface protein expression was assessed by labeling the flanking tags (c-Myc and hemagglutinin) via flow cytometry. C-Myc was labeled with 1:1000 chicken anti-cMyc (Exalpha) followed by 1:1000 goat anti-chicken AF488 (Invitrogen). Hemagglutinin was labeled with 1:1000 mouse anti-hemagglutinin (clone 16B12; Biolegend) followed by 1:1000 goat anti-mouse AF647 (Invitrogen). Binding was assessed by incubating yeast with 1 µM of 250 kDa monobiotinylated hyaluronic acid (CD Bioparticles) for 1 hour, followed by secondary labeling with 1:1000 streptavidin-AF647 (Invitrogen).

Following identification of versican G1 as the lead HABP, mutagenesis of the wildtype sequence was performed using error-prone PCR. The resulting DNA library was transformed into yeast using two-piece homologous recombination and electroporation. The resulting yeast library was grown and induced as described above. Magnetic bead selection followed by FACS was used to isolate improved binders. Once library convergence was achieved, single clones were isolated for sequencing and aligned to the wildtype sequence. Once a single clone of our improved mutant was isolated, yeast encoding the wildtype or mutant sequence were grown at 30°C in SD-CAA and induced at either 20°C, 30°C, or 37°C in SG-CAA before titration with serial dilutions of 250 kDa monobiotinylated HA to generate binding curves. Yeast were sorted on a FACS Aria III Cell Sorter (BD) and analyzed using either FACS Symphony A3 (BD), FACS Symphony A1 (BD), or FACS LSR Fortessa (BD)

### Bio-layer interferometry

Bio-layer interferometry (BLI) was performed using an Octet RED96e instrument (ForteBio, Sartorius AG). PBS supplemented with 0.1% (w/v) BSA and 0.05% (v/v) Tween 20 (Sigma Aldrich) was used as buffer in this experiment. 40 nM of 250 kDa monobiotinylated hyaluronic acid was loaded onto streptavidin sensors (ForteBio, Sartorius AG) for 60 seconds. Sensors were then moved into buffer for 60 seconds to establish a baseline, and subsequently moved into serial dilutions of protein. Protein was associated for 500 seconds before sensors were moved to buffer to track protein disassociation. The protein concentrations used in all BLI experiments were 2000 nM, 1000 nM, 500 nM, 250 nM, 125 nM, and 62.5 nM. Kinetic parameters were fit using Octet System Data Analysis software (ForteBio, Sartorius AG).

### Collagen ELISA

Collagen I coated 96-well plates (Gibco) were blocked in PBS supplemented with 0.1% (w/v) BSA and 0.05% (v/v) Tween 20 for 1 hour at room temperature. This buffer was used for the entirety of the experiment. Wells were titrated with the indicated concentrations of lumican fusions for 2 hours at room temperature. Wells were washed and then incubated for 1 hour with horseradish peroxidase-conjugated polyclonal anti-6xHis (Abcam) at 1:4000 dilution or polyclonal horseradish peroxidase-conjugated anti-mouse IgG H&L (Abcam) at 1:2000 dilution. Wells were washed again, and then 1-Step Ultra TMB-ELISA Substrate Solution (Thermo Fisher Scientific) was added for 5 minutes. 1M sulfuric acid was then added to stop the reaction. Data acquisition occurred on Synergy HTX Multimode plate reader (BioTek) (absorbance at 450 nm, corrected with a reference absorbance at 570 nm).

### Thermal shift assay

The protein thermal shift assay was performed using SYPRO Orange dye (Thermo Fisher Scientific), following vendors instructions. Briefly, 2.5 µL of 50x SYPRO Orange was added to 10 µg of purified protein. The reaction mixture was adjusted to a final volume of 25 µL with PBS before melting and differential scanning fluorimetry was performed on a LightCycler 480 II (Roche).

### Cytokine activity assays

For IL-12 bioactivity, activity was assessed using the HEK-Blue IL-12 reporter assay (Invivogen) according to manufacturer’s instructions. For IL-15 bioactivity, CTLL-2 cells were cultured according to vendor instructions before they were starved of IL-2 24 hours prior to exposure to serial dilution of IL-15 fusion proteins. Cells were incubated for 2 days before addition of CellTiterGlo 2.0 (Promega) to assess cell number via ATP levels. Data acquisition for these assays was performed on Synergy HTX Multimode plate reader.

### Tumor inoculation and treatment

All mice received tumor inoculums of one million cells in 50 µL of sterile PBS. Tumor cells were injected onto the shaved right flank of mice. Prior to treatment, tumors were measured and mice were binned into groups to ensure equal mean tumor size between study groups. Caliper measurements of tumor length and width were used to determine tumor area. All proteins were diluted in sterile PBS before intratumoral administration of 20 µL injections. For efficacy studies, mice were treated on days 6 and 13 after tumor inoculation. Mice received 0.11 nmol IL-15 (IgG-15, 10.6 µg; lumican-Fc-IL15, 9.5 µg; versican S139G_T_-Fc-IL15, 9.3 µg; 1B7-1B7-MSA-IL15, 13 µg; lumican-MSA-IL15, 13.9 µg; versican S139G_T_-MSA-IL15, 13.8 µg) and 0.014 nmol IL-12 (IL12-IgG, 1.9 µg; IL12-lumican-Fc, 1.7 µg; IL12-versican S139G_T_-Fc, 1.7 µg; IL12-1B7-1B7-MSA, 2.2 µg; IL12-lumican-MSA, 2.3 µg; IL12-versican S139G_T_-MSA, 2.3 µg). Mice were euthanized when tumor area exceeded 100 mm^2^ or if weight loss exceeded 20%. For toxicity experiments, mice were treated 6 days after tumor inoculation. Mice received 0.22 nmol IL-15 (1B7-1B7-MSA-IL15, 25.9 µg; lumican-MSA-IL15, 27.9 µg; versican S139G_T_-MSA-IL15, 27.5 µg) and 0.028 nmol IL-12 (IL12-1B7-1B7-MSA, 4.4 µg; IL12-lumican-MSA, 4.6 µg; IL12-versican S139G_T_-MSA, 4.5 µg). Mice were euthanized if weight loss exceeded 20%. For the histology study, mice were treated on days 6 and 13 with 0.11 nmol IL-15 (IgG-15, 10.6 µg; lumican-Fc-IL15, 9.5 µg; versican S139G_T_-Fc-IL15, 9.3 µg) and 0.014 nmol IL-12 (IL12-IgG, 1.9 µg; IL12-lumican-Fc, 1.7 µg; IL12-versican S139G_T_-Fc, 1.7 µg). For *in vivo* pSTAT4 assessment, mice were treated 8 days after tumor induction with 0.06 nmol IL-12 (IL12-1B7-1B7-MSA, 9.4 µg; IL12-lumican-MSA, 9.9 µg; IL12-versican S139G_T_-MSA, 9.9 µg)

### IVIS imaging

Proteins were labeled with AlexaFluor 647 NHS Ester (Invitrogen) according to manufacturer’s instructions. Excess dye was removed using Zeba 7kDa MWCO desalting columns (Thermo Fisher Scientific). Degree of dye labeling was assessed using a NanoDrop (Thermo Fisher Scientific). 0.1 nmol each of control unanchored IgG (14.8 µg), lumican-Fc (13.2 µg), and versican S139G_T_-Fc (12.7 µg) were intratumorally injected into B16F10-TRP2 KO tumor bearing albino mice (B6(Cg)-Tyr^c-2J^/J) 7 days after tumor inoculation. All doses contained equimolar dye. Following injection, mice were imaged with the IVIS Spectrum Imaging System 100 (IVIS; Xenogen) for 9 days. Mice were fed an alfalfa-free diet (Test Diet) throughout the experiment, starting 3 days prior to imaging. Tumor-localized total radiant efficiency was obtained via data analysis on Living Image software (Caliper Life Sciences).

### Biodistribution

Proteins were fluorescently labeled as previously described. Proteins were administered at indicated doses intratumorally into B16F10 or MC38 tumor-bearing mice 7 days after tumor induction. Twenty-four hours after dosing, tumor and tumor draining lymph nodes were dissected, and blood was collected via intracardiac puncture. Serum was isolated from blood using serum gel tubes (SAI Infusion Technologies) according to manufacturers instructions. Tumors were weighed before resuspension in Pierce IP Lysis Buffer (Thermo Fisher Scientific) supplemented with Pierce Protease Inhibitor tablet (Thermo Fisher Scientific) at 15 µL buffer/mg tumor prior to homogenization with gentleMACS M tubes on a gentleMACS Octo-dissociator (Miltenyi). Lymph nodes were weighed prior to manual homogenization in 150 µL lysis buffer with protease inhibitor. Compartment fluorescence was assessed using a Synergy HTX Multimode plate reader, and protein concentration calculated via a standard curve.

### Immunofluorescent microscopy

Proteins were fluorescently labeled as previously described and injected intratumorally into established B16F10 tumors 7 days after tumor induction. Twenty-four hours after protein injection, tumors were excised and fixed overnight in 4% paraformaldehyde (PFA; Invitrogen). Tumors were cryoprotected by sequential incubation in 15% and 30% sucrose solutions (Thermo Fisher Scientific), each overnight. Following sucrose infiltration, tumors were embedded in optimal cutting temperature compound (OCT; Scigen), frozen, and sectioned to 10 µm on a cryostat (Leica Biosystems). Sections were stored at -20°C until use. Prior to staining, sections were dried at room temperature and washed with PBS. Nuclear staining was performed with 0.1 µg/mL DAPI (Invitrogen) at room temperature for 5 minutes and subsequently washed three times with PBS. Sections were mounted with ProLong Diamond Antifade Mountant (Thermo Fisher Scientific) before imaging using a TissueFAXs SL Fluorescent slide scanner (TissueGnostics) equipped with a 25X air objective. All image analysis was performed in Fiji. Tumor area was defined as DAPI-positive regions. For each section, an intensity threshold was set based on average and standard deviation of pixel intensity of uninjected, DAPI-stained tumors. Percent tumor coverage was calculated as the fraction of pixels exceeding this threshold relative to the total number of tumor pixels. Integrated intensity was calculated by summing all pixel intensities within the tumor area and subtracting background values measured from blank B16F10 tumor sections.

### Legendplex and ALT analysis

B16F10 tumor bearing mice were treated as described with IL-15/IL-12 combination therapy. Blood was collected by submandibular bleeding 1- and 3-days post-treatment. Mice were bled directly into serum gel collection tubes. Tubes were centrifuged at 10,000g for 10 minutes before serum was isolated and stored at -80°C until use. The colorimetric alanine transaminase activity assay kit (Abcam) was used to determine circulating ALT levels. Samples were diluted 1:5 and vendor instructions were followed. Sample plates were read using a Synergy HTX Multimode plate reader. For cytokine/chemokine analysis, the Legendplex Mouse Cytokine Release Syndrome panel (Biolegend) was used according to vendor instructions. Serum samples were diluted 1:2 or 1:4, and sample plates were run on a FACS LSR Fortessa (BD). Data analysis was performed using Biolegend’s LEGENDplex data analysis software.

### Histological assessment

Mice bearing B16F10 tumors were treated 6 and 13 days after tumor inoculation with IL-12/IL-15 cytokine therapy intratumorally and subcutaneously on contralateral healthy flank. Half the mice were euthanized on day 13 before therapy administration, and the other half on day 19, 6 days after their last treatment. Once euthanized, tumors and skin from the treated contralateral flank were dissected and placed in 10% formalin (Sigma Aldrich) overnight. Tissues were then mounted in paraffin, sectioned to 5 µm, and H&E stained for histopathological analysis. Slides were imaged using an Aperio Digital Slide Scanner at 20X. All samples were evaluated by a board-certified veterinary pathologist.

### pSTAT flow cytometry

*In vivo* assessment of STAT phosphorylation was performed as previously described (*44*). Briefly, B16F10 tumors were weighed before processing into single-cell suspensions through a 70-µm strainer directly into PBS supplemented with 1:1000 Zombie Live Dead NIR (Biolegend) with Halt Protease and 1x Phosphatase Inhibitor Cocktail (Thermo Fisher Scientific). Samples were incubated at room temperature for 5 minutes before they were washed and subsequently fixed with BD Phosflow Fixation Buffer I, prewarmed to 37°C. Samples were incubated at room temperature for 7 minutes before washing and addition of 25 mg to the assay plate. At this stage, 10,000 precision counting beads (Biolegend) were added to each sample. Samples were subsequently permeabilized with BD Phosflow Perm Buffer III which was prechilled to -20°C. Following a 30 minutes incubation on ice, samples were stained with 1:100 anti-pSTAT4 PE (clone 38; BD Biosciences), 1:100 anti-CD3 BUV395 (clone 17A2; Invitrogen), and 1:100 anti-CD45 BUV496 (clone 30-F11; BD Biosciences). Samples were stored in the dark at 4°C overnight. Samples were washed before data acquisition using a BD FACSymphony A3. Data were analyzed in FlowJo.

### Statistical analysis

Analyses were performed using GraphPad Prism 10 software. Comparisons between groups were assessed by one-way or two-way ANOVA with Tukey’s multiple comparisons test. Survival analysis was done using log-rank Mantel-Cox test. Significance is as follows: ns, not significant; *P < 0.05; **P < 0.01; ***P<0.001; ****P < 0.0001. Study power and statistical methods are present in figure legends. Raw data is presented in data files S1 and S2.

## ACKKNOWLEDGEMENTS

We thank the staff of the Division for Comparative Medicine at MIT for support in animal care during *in vivo* studies. We additionally thank the staff of the Koch Institute Swanson Biotechnology Center (National Cancer Institute Grant P30-CA14051) for technical assistance. Special thanks to Dr. Magalie Boucher, DVM, MS, ACVP for the histopathology evaluation and pathology expertise.

## FUNDING

National Science Foundation Graduate Fellowship program (WPIII)

George and Daisy Soros Fellowship for New Americans (RA)

Ludwig Center Koch Institute Graduate Fellowship (LD)

## AUTHOR CONTRIBUTIONS

E.F. and K.D.W. conceived, designed, and directed this study. E.F., W.P.III, L.D., R.A., D.K., and J.S. performed experiments. E.F. performed data analysis. E.F. and K.D.W. wrote the manuscript, with input from all authors.

## COMPETING INTERESTS

K.D.W is a named inventor on U.S. Provisional Patent no. US20200102370A1, *Collagen-localized immunomodulatory molecules and methods thereof*. K.D.W. and E.F. are named inventors on U.S. Provisional Patent application no. 64/007,109, *Hyaluronic acid anchor immunotherapies and methods thereof*. All other authors declare they have no competing interests.

## REFERENCES

1. S. Paul, M. F. Konig, D. M. Pardoll, C. Bettegowda, N. Papadopoulos, K. M. Wright, S. B. Gabelli, M. Ho, A. van Elsas, S. Zhou, Cancer therapy with antibodies. Nat. Rev. Cancer 24, 399–426 (2024).

2. D. J. Propper, F. R. Balkwill, Harnessing cytokines and chemokines for cancer therapy. Nat. Rev. Clin. Oncol. 19, 237–253 (2022).

3. M. Srinivasarao, C. V. Galliford, P. S. Low, Principles in the design of ligand-targeted cancer therapeutics and imaging agents. Nat. Rev. Drug Discov. 14, 203–219 (2015).

4. L. Santollani, K. D. Wittrup, Spatiotemporally programming cytokine immunotherapies through protein engineering. Immunol. Rev. 320, 10–28 (2023).

5. M. Li, S. Mei, Y. Yang, Y. Shen, L. Chen, Strategies to mitigate the on- and off-target toxicities of recombinant immunotoxins: an antibody engineering perspective. Antib. Ther. 5, 163–175 (2022).

6. C. M. V. Herpen, R. Huijbens, M. Looman, J. D. Vries, H. Marres, J. V. D. Ven, R. Hermsen, G. J. Adema, P. H. D. Mulder, Pharmacokinetics and immunological aspects of a phase Ib study with intratumoral administration of recombinant human interleukin-12 in patients with head and neck squamous cell carcinoma: a decrease of T-bet in peripheral blood mononuclear cells. Clin Cancer Res Official J Am Assoc Cancer Res 9, 2950–6 (2003).

7. Y. Agarwal, L. E. Milling, J. Y. H. Chang, L. Santollani, A. Sheen, E. A. Lutz, A. Tabet, J. Stinson, K. Ni, K. A. Rodrigues, T. J. Moyer, M. B. Melo, D. J. Irvine, K. D. Wittrup, Intratumourally injected alum-tethered cytokines elicit potent and safer local and systemic anticancer immunity. Nat Biomed Eng 6, 129–143 (2022).

8. A. R. Chakravarti, C. E. Groer, H. Gong, V. Yudistyra, M. L. Forrest, C. J. Berkland, Design of a Tumor Binding GMCSF as Intratumoral Immunotherapy of Solid Tumors. Mol. Pharm. 20, 1975–1989 (2023).

9. R. Danielli, R. Patuzzo, A. M. D. Giacomo, G. Gallino, A. Maurichi, A. D. Florio, O. Cutaia, A. Lazzeri, C. Fazio, C. Miracco, L. Giovannoni, G. Elia, D. Neri, M. Maio, M. Santinami, Intralesional administration of L19-IL2/L19-TNF in stage III or stage IVM1a melanoma patients: results of a phase II study. *Cancer Immunol.*, Immunother. 64, 999–1009 (2015).

10. N. Momin, N. K. Mehta, N. R. Bennett, L. Ma, J. R. Palmeri, M. M. Chinn, E. A. Lutz, B. Kang, D. J. Irvine, S. Spranger, K. D. Wittrup, Anchoring of intratumorally administered cytokines to collagen safely potentiates systemic cancer immunotherapy. Sci Transl Med 11 (2019), doi:10.1126/scitranslmed.aaw2614.

11. J. R. Palmeri, B. M. Lax, J. M. Peters, L. Duhamel, J. A. Stinson, L. Santollani, E. A. Lutz, W. Pinney, B. D. Bryson, K. D. Wittrup, CD8+ T cell priming that is required for curative intratumorally anchored anti-4-1BB immunotherapy is constrained by Tregs. Nat. Commun. 15, 1900 (2024).

12. E. A. Lutz, N. Jailkhani, N. Momin, Y. Huang, A. Sheen, B. H. Kang, K. D. Wittrup, R. O. Hynes, Intratumoral nanobody–IL-2 fusions that bind the tumor extracellular matrix suppress solid tumor growth in mice. PNAS Nexus 1, pgac244 (2022).

13. P. Chen, B. M. Bordeau, W. Zhang, J. P. Balthasar, Investigations of Influence of Antibody Binding Kinetics on Tumor Distribution and Anti-Tumor Efficacy. AAPS J. 27, 91 (2025).

14. N. Momin, J. R. Palmeri, E. A. Lutz, N. Jailkhani, H. Mak, A. Tabet, M. M. Chinn, B. H. Kang, V. Spanoudaki, R. O. Hynes, K. D. Wittrup, Maximizing response to intratumoral immunotherapy in mice by tuning local retention. Nat Commun 13, 109 (2022).

15. T. R. Cox, The matrix in cancer. Nat. Rev. Cancer 21, 217–238 (2021).

16. L. Arpinati, G. Carradori, R. Scherz-Shouval, CAF-induced physical constraints controlling T cell state and localization in solid tumours. Nat. Rev. Cancer 24, 676–693 (2024).

17. J. R. E. Fraser, T. C. Laurent, U. B. G. Laurent, Hyaluronan: its nature, distribution, functions and turnover. J. Intern. Med. 242, 27–33 (1997).

18. R. K. Boregowda, H. N. Appaiah, M. Siddaiah, S. B. Kumarswamy, S. Sunila, T. KN, K. Mortha, B. Toole, S. d Banerjee, Expression of Hyaluronan in human tumor progression. J Carcinog 5, 2–2 (2006).

19. G. Alexandrakis, E. B. Brown, R. T. Tong, T. D. McKee, R. B. Campbell, Y. Boucher, R. K. Jain, Two-photon fluorescence correlation microscopy reveals the two-phase nature of transport in tumors. Nat Med 10, 203–207 (2004).

20. M. Liu, C. Tolg, E. Turley, Dissecting the Dual Nature of Hyaluronan in the Tumor Microenvironment. Front. Immunol. 10, 947 (2019).

21. J. Winkler, A. Abisoye-Ogunniyan, K. J. Metcalf, Z. Werb, Concepts of extracellular matrix remodelling in tumour progression and metastasis. Nat. Commun. 11, 5120 (2020).

22. J. I. Park, L. Cao, V. M. Platt, Z. Huang, R. A. Stull, E. E. Dy, J. J. Sperinde, J. S. Yokoyama, F. C. Szoka, Antitumor Therapy Mediated by 5-Fluorocytosine and a Recombinant Fusion Protein Containing TSG-6 Hyaluronan Binding Domain and Yeast Cytosine Deaminase. Mol Pharmaceut 6, 801–812 (2009).

23. J. Liu, D. Pan, X. Huang, S. Wang, H. Chen, Y. Z. Zhu, L. Ye, Targeting collagen in tumor extracellular matrix as a novel targeted strategy in cancer immunotherapy. Front. Oncol. 13, 1225483 (2023).

24. H. Ikemoto, P. Lingasamy, A.-M. A. Willmore, H. Hunt, K. Kurm, O. Tammik, P. Scodeller, L. Simón-Gracia, V. R. Kotamraju, A. M. Lowy, K. N. Sugahara, T. Teesalu, Hyaluronan-binding peptide for targeting peritoneal carcinomatosis. Tumor Biol. 39, 1010428317701628 (2017).

25. J. K. Cini, S. Dexter, D. J. Rezac, S. J. McAndrew, G. Hedou, R. Brody, R.-N. Eraslan, R. T. Kenney, P. Mohan, SON-1210 - a novel bifunctional IL-12 / IL-15 fusion protein that improves cytokine half-life, targets tumors, and enhances therapeutic efficacy. Front. Immunol. 14, 1326927 (2023).

26. K. Matsumoto, M. Shionyu, M. Go, K. Shimizu, T. Shinomura, K. Kimata, H. Watanabe, Distinct Interaction of Versican/PG-M with Hyaluronan and Link Protein*. J. Biol. Chem. 278, 41205–41212 (2003).

27. J. Lesley, N. M. English, I. Gál, K. Mikecz, A. J. Day, R. Hyman, Hyaluronan Binding Properties of a CD44 Chimera Containing the Link Module of TSG-6*. J. Biol. Chem. 277, 26600–26608 (2002).

28. R. Peach, D. Hollenbaugh, I. Stamenkovic, A. Aruffo, Identification of hyaluronic acid binding sites in the extracellular domain of CD44. J. cell Biol. 122, 257–264 (1993).

29. J. Jumper, R. Evans, A. Pritzel, T. Green, M. Figurnov, O. Ronneberger, K. Tunyasuvunakool, R. Bates, A. Žídek, A. Potapenko, A. Bridgland, C. Meyer, S. A. A. Kohl, A. J. Ballard, A. Cowie, B. Romera-Paredes, S. Nikolov, R. Jain, J. Adler, T. Back, S. Petersen, D. Reiman, E. Clancy, M. Zielinski, M. Steinegger, M. Pacholska, T. Berghammer, S. Bodenstein, D. Silver, O. Vinyals, A. W. Senior, K. Kavukcuoglu, P. Kohli, D. Hassabis, Highly accurate protein structure prediction with AlphaFold. Nature 596, 583–589 (2021).

30. Y. J. Wu, D. P. L. Pierre, J. Wu, A. J. Yee, B. B. Yang, The interaction of versican with its binding partners. Cell Res. 15, 483–494 (2005).

31. M. Lo, H. S. Kim, R. K. Tong, T. W. Bainbridge, J.-M. Vernes, Y. Zhang, Y. L. Lin, S. Chung, M. S. Dennis, Y. J. Y. Zuchero, R. J. Watts, J. A. Couch, Y. G. Meng, J. K. Atwal, R. J. Brezski, C. Spiess, J. A. Ernst, Effector-attenuating Substitutions That Maintain Antibody Stability and Reduce Toxicity in Mice*. J. Biol. Chem. 292, 3900–3908 (2017).

32. M. Sternke, K. W. Tripp, D. Barrick, Consensus sequence design as a general strategy to create hyperstable, biologically active proteins. Proc National Acad Sci 116, 11275–11284 (2019).

33. M. Fang, J. Yuan, C. Peng, Y. Li, Collagen as a double-edged sword in tumor progression. Tumor Biol. 35, 2871–2882 (2014).

34. J. Riegler, Y. Labyed, S. Rosenzweig, V. Javinal, A. Castiglioni, C. X. Dominguez, J. E. Long, Q. Li, W. Sandoval, M. R. Junttila, S. J. Turley, J. Schartner, R. A. D. Carano, Tumor Elastography and Its Association with Collagen and the Tumor Microenvironment. Clin Cancer Res 24, 4455–4467 (2018).

35. L. Santollani, L. Maiorino, Y. J. Zhang, J. R. Palmeri, J. A. Stinson, L. R. Duhamel, K. Qureshi, J. R. Suggs, O. T. Porth, W. Pinney, R. A. Msari, A. A. Walsh, K. D. Wittrup, D. J. Irvine, Local delivery of cell surface-targeted immunocytokines programs systemic antitumor immunity. Nat. Immunol. 25, 1820–1829 (2024).

36. S. I. S. Mosely, J. E. Prime, R. C. A. Sainson, J.-O. Koopmann, D. Y. Q. Wang, D. M. Greenawalt, M. J. Ahdesmaki, R. Leyland, S. Mullins, L. Pacelli, D. Marcus, J. Anderton, A. Watkins, J. C. Ulrichsen, P. Brohawn, B. W. Higgs, M. McCourt, H. Jones, J. A. Harper, M. Morrow, V. Valge-Archer, R. Stewart, S. J. Dovedi, R. W. Wilkinson, Rational Selection of Syngeneic Preclinical Tumor Models for Immunotherapeutic Drug Discovery. Cancer Immunol Res 5, 29–41 (2017).

37. R. Afik, E. Zigmond, M. Vugman, M. Klepfish, E. Shimshoni, M. Pasmanik-Chor, A. Shenoy, E. Bassat, Z. Halpern, T. Geiger, I. Sagi, C. Varol, Tumor macrophages are pivotal constructors of tumor collagenous matrix. J. Exp. Med. 213, 2315–2331 (2016).

38. K. Pfisterer, L. E. Shaw, D. Symmank, W. Weninger, The Extracellular Matrix in Skin Inflammation and Infection. Front. Cell Dev. Biol. 9, 682414 (2021).

39. H. S. Abyaneh, M. Regenold, T. D. McKee, C. Allen, M. A. Gauthier, Towards extracellular matrix normalization for improved treatment of solid tumors. Theranostics 10, 1960–1980 (2020).

40. K. M. Mak, R. Mei, Basement Membrane Type IV Collagen and Laminin: An Overview of Their Biology and Value as Fibrosis Biomarkers of Liver Disease. Anat. Rec. 300, 1371–1390 (2017).

41. K. D. Moynihan, C. F. Opel, G. L. Szeto, A. Tzeng, E. F. Zhu, J. M. Engreitz, R. T. Williams, K. Rakhra, M. H. Zhang, A. M. Rothschilds, S. Kumari, R. L. Kelly, B. H. Kwan, W. Abraham, K. Hu, N. K. Mehta, M. J. Kauke, H. Suh, J. R. Cochran, D. A. Lauffenburger, K. D. Wittrup, D. J. Irvine, Eradication of large established tumors in mice by combination immunotherapy that engages innate and adaptive immune responses. Nat. Med. 22, 1402–1410 (2016).

42. J. Zhou, G. G. Yan, D. Cluckey, C. Meade, M. Ruth, R. Sorm, A. S. Tam, S. Lim, C. Petridis, L. Lin, A. M. D’Antona, X. Zhong, Exploring Parametric and Mechanistic Differences between Expi293FTM and ExpiCHO-STM Cells for Transient Antibody Production Optimization. Antibodies 12, 53 (2023).

43. B. H. Kang, B. M. Lax, K. D. Wittrup, Yeast Surface Display M. W. Traxlmayr, Ed. (2022).

44. S. Gaggero, J. Martinez-Fabregas, A. Cozzani, P. K. Fyfe, M. Leprohon, J. Yang, F. E. Thomasen, H. Winkelmann, R. Magnez, A. G. Conti, S. Wilmes, E. Pohler, M. van G. Bonnello, X. Thuru, B. Quesnel, F. Soncin, J. Piehler, K. Lindorff-Larsen, R. Roychoudhuri, I. Moraga, S. Mitra, IL-2 is inactivated by the acidic pH environment of tumors enabling engineering of a pH-selective mutein. Sci. Immunol. 7, eade5686 (2022).

