## Supplementary Tables and Figures for "Engineering hyaluronic acid-binding cytokines for enhanced tumor retention and safety"

**Supplementary Table S1**

Kinetic binding parameters of wildtype and mutant versican proteins, as determined by BLI.

| Construct | $K_{D,1}$ [M] | $K_{D,2}$ [M] | $k_{a,1}$ [$M^{-1}s^{-1}$] | $k_{a,2}$ [$M^{-1}s^{-1}$] | $k_{off,1}$ [$s^{-1}$] | $k_{off,2}$ [$s^{-1}$] |
|---|---|---|---|---|---|---|
| Vcan MSA | 0.009 | - | 3.05 ± 0.126 | - | 0.027 ± 1.19 e-3 | - |
| Vcan S139G MSA | 8.19 e-7 | - | 9.14 e3 ± 114 | - | 7.48 e-3 ± 5.90 e-5 | - |
| Vcan S139G$_T$ MSA | 1.06 e-6 | - | 7.87 e3 ± 87 | - | 8.33 e-3 ± 5.50e-5 | - |
| Vcan Fc | 4.38 e-8 | 1.65 e-7 | 5.99 e3 ± 17.8 | 1.24 e5 ± 1.28 e3 | 2.62 e-4 ± 1.69 e-4 | 0.024 ± 9.48 e-5 |
| Vcan S139G Fc | 1.05 e-11 | 4.86 e-8 | 1.08 e4 ± 35.0 | 1.22 e5 ± 780 | 1.14 e-7 ± 2.09 e-7 | 5.91 e-3 ± 1.76 e-5 |
| Vcan S139G$_T$ Fc | 4.19 e-9 | 4.87 e-8 | 1.14 e4 ± 53.4 | 1.38 e5 ± 1.31 e3 | 4.76 e-5 ± 5.02 e-6 | 6.71 e-3 ± 3.34e-5 |

**Supplementary Table S2**

Amino acid sequences of proteins of yeast-displayed and recombinantly produced proteins.

Key: Targeted or untargeted protein, VHH, or $V_H/V_L$, Linker, Fc, MSA, Cytokine, His tag

| | |
|---|---|
| CD44 | QIDLNVTCRYAGVFHVEKNGRYSISRTEAADLCQAFNSTLPTMDQMKLALSKGFETCRYGFIEGNVVIPRIHPNAICAANHTGVYILVTSNTSHYDTYCFNASAPPEEDCTSVTDLPNSFDGPVTITIVNRDGTRYSKKGEYRTHQEDIDASNIIDDDVSSGSTIEKSTPEGYILHTYLPTEQPTGDQDDSFFIRSTLAT |
| CD44-TSG Link Chimera | QIDLNVTCRYAGVYHREARAGRYKLTYAEAKAVCEFEGGRLATYKQLEAARKIGFHVCAAGWMAKGRVGYPIVKPGPNCGFGKTGIIDYGIRLNRSERWDAYCYNASAPPEEDCTSVTDLPNSFDGPVTITIVNRDGTRYSKKGEYRTHQEDIDASNIIDDDVSSGSTIEKSTPEGYILHTYLPTEQPTGDQDDSFFIRSTLAT |
| HAPLN1 | MRSLLLLVLISVCWADHHLSDSYTPPDQDRVIHIQAENGPRLLVEAEQAKVFSHRGGNVTLPCKFYRDPTAFGSGIHKIRIKWTKLTSDYLREVDVFVSMGYHKKTYGGYQGRVFLKGGSDNDASLVITDLTLEDYGRYKCEVIEGLEDDTAVVALELQGVVFPYFPRLGRYNLNFHEARQACLDQDAVIASFDQLYDAWRGGLDWCNAGWLSDGSVQYPITKPREPCGGQNTVPGVRNYGFWDKDKSRYDVFCFTSNFNGRFYYLIHPTKLTYDEAVQACLNDGAQIAKVGQIFAAWKLLGYDRCDAGWLADGSVRYPISRPRRRCSPTEAAVRFVGFPDKKHKLYGVYCFRAYN |
| Versican (S139 position highlighted) | LHQAKMETSPPVKGSLSGKVVLPCHFSTLPTLPPNYNTSEFLRIKWSKMEVDKNGKDIKETTVLVAQNGNIKIGQDYKGRVSVPTHPDDVGDASLTMVKLRASDAAVYRCDVMYGIEDTQDTMSLAVDGVVFHYRAATSRYTLNFAAAQQACLDIGAVIASPEQLFAAYEDGFEQCDAGWLSDQTVRYPIRAPREGCYGDMMGKEGVRTYGFRSPQETYDVYCYVDHLDGDVFHITAPSKFTFEEAEAECTSRDARLATVGELQAAWRNGFDQCDYGWLSDASVRHPVTVARAQCGGGLLGVRTLYRFENQTCFPLPDSRFDAYCF |
| Versican-MSA | LHQAKMETSPPVKGSLSGKVVLPCHFSTLPTLPPNYNTSEFLRIKWSKMEVDKNGKDIKETTVLVAQNGNIKIGQDYKGRVSVPTHPDDVGDASLTMVKLRASDAAVYRCDVMYGIEDTQDTMSLAVDGVVFHYRAATSRYTLNFAAAQQACLDIGAVIASPEQLFAAYEDGFEQCDAGWLSDQTVRYPIRAPREGCYGDMMGKEGVRTYGFRSPQETYDVYCYVDHLDGDVFHITAPSKFTFEEAEAECTSRDARLATVGELQAAWRNGFDQCDYGWLSDASVRHPVTVARAQCGGGLLGVRTLYRFENQTCFPLPDSRFDAYCFGGGSGGGSEAHKSEIAHRYNDLGEQHFKGLVLIAFSQYLQKCSYDEHAKLVQEVTDFAKTCVADESAANCDKSLHTLFGDKLCAIPNLRENYGELADCCTKQEPERNECFLQHKDDNPSLPPFERPEAEAMCTSFKENPTTFMGHYLHEVARRHPYFYAPELLYYAEQYNEILTQCCAEADKESCLTPKLDGVKEKALVSSVRQRMKCSSMQKFGERAFKAWAVARLSQTFPNADFAEITKLATDLTKVNKECCHGDLLECADDRAELAKYMCENQATISSKLQTCCDKPLLKKAHCLSEVEHDTMPADLPAIAADFVEDQEVCKNYAEAKDVFLGTFLYEYSRRHPDYSVSLLLRLAKKYEATLEKCCAEANPPACYGTVLAEFQPLVEEPKNLVKTNCDLYEKLGEYGFQNAILVRYTQKAPQVSTPTLVEAARNLGRVGTKCCTLPEDQRLPCVEDYLSAILNRVCLLHEKTPVSEHVTKCCSGSLVERRPCFSALTVDETYVPKEFKAETFTFHSDICTLPEKEKQIKKQTALAELVKHKPKATAEQLKTVMDDFAQFLDTCCKAADKDTCFSTEGPNLVTRCKDALAHHHHHH |
| Versican-Fc (mIgG2C LALA-PG) | LHQAKMETSPPVKGSLSGKVVLPCHFSTLPTLPPNYNTSEFLRIKWSKMEVDKNGKDIKETTVLVAQNGNIKIGQDYKGRVSVPTHPDDVGDASLTMVKLRASDAAVYRCDVMYGIEDTQDTMSLAVDGVVFHYRAATSRYTLNFAAAQQACLDIGAVIASPEQLFAAYEDGFEQCDAGWLSDQTVRYPIRAPREGCYGDMMGKEGVRTYGFRSPQETYDVYCYVDHLDGDVFHITAPSKFTFEEAEAECTSRDARLATVGELQAAWRNGFDQCDYGWLSDASVRHPVTVARAQCGGGLLGVRTLYRFENQTCFPLPDSRFDAYCFGGGGSGGGGSGGGGSEPRVPITQNPCPPLKECPPCAAPDAAGGPSVFIFPPKIKDVLMISLSPMVTCVVVDVSEDDPDVQISWFVNNVEVHTAQTQTHREDYNSTLRVVSALPIQHQDWMSGKEFKCKVNNRALGSPIEKTISKPRGPVRAPQVYVLPPPAEEMTKKEFSLTCMITGFLPAEIAVDWTSNGRTEQNYKNTATVLDSDGSYFMYSKLRVQKSTWERGSLFACSVVHEGLHNHLTTKTISRSLGK |
| Versican S139G-MSA | LHQAKMETSPPVKGSLSGKVVLPCHFSTLPTLPPNYNTSEFLRIKWSKMEVDKNGKDIKETTVLVAQNGNIKIGQDYKGRVSVPTHPDDVGDASLTMVKLRASDAAVYRCDVMYGIEDTQDTMSLAVDGVVFHYRAATGRYTLNFAAAQQACLDIGAVIASPEQLFAAYEDGFEQCDAGWLSDQTVRYPIRAPREGCYGDMMGKEGVRTYGFRSPQETYDVYCYVDHLDGDVFHITAPSKFTFEEAEAECTSRDARLATVGELQAAWRNGFDQCDYGWLSDASVRHPVTVARAQCGGGLLGVRTLYRFENQTCFPLPDSRFDAYCFGGGSGGGSEAHKSEIAHRYNDLGEQHFKGLVLIAFSQYLQKCSYDEHAKLVQEVTDFAKTCVADESAANCDKSLHTLFGDKLCAIPNLRENYGELADCCTKQEPERNECFLQHKDDNPSLPPFERPEAEAMCTSFKENPTTFMGHYLHEVARRHPYFYAPELLYYAEQYNEILTQCCAEADKESCLTPKLDGVKEKALVSSV |

| | |
|---|---|
| | RQRMKCSSMQKFGERAFKAWAVARLSQTFPNADFAEITKLATDLTKVNKECCHGDLLECADDRAEL<br>AKYMCENQATISSKLQTCCDKPLLKKAHCLSEVEHDTMPADLPAIAADFVEDQEVCKNYAEAKDVFL<br>GTFLYEYSRRHPDYSVSLLLRLAKKYEATLEKCCAEANPPACYGTVLAEFQPLVEEPKNLVKTNCDL<br>YEKLGEYGFQNAILVRYTQKAPQVSTPTLVEAARNLGRVGTKCCTLPEDQRLPCVEDYLSAILNRVC<br>LLHEKTPVSEHVTKCCSGSLVERRPCFSALTVDETYVPKEFKAETFTFHSDICTLPEKEKQIKKQTAL<br>AELVKHKPKATAEQLKTVMDDFAQFLDTCCKAADKDTCFSTEGPNLVTRCKDALAHHHHHH |
| Versican S139G-Fc (mIgG2C LALA-PG) | LHQAKMETSPPVKGSLSGKVVLPCHFSTLPTLPPNYNTSEFLRIKWSKMEVDKNGKDIKETTVLVAQ<br>NGNIKIGQDYKGRVSVPTHPDDVGDASLTMVKLRASDAAVYRCDVMYGIEDTQDTMSLAVDGVVFH<br>YRAATGRYTLNFAAAQQACLDIGAVIASPEQLFAAYEDGFEQCDAGWLSDQTVRYPIRAPREGCYG<br>DMMGKEGVRTYGFRSPQETYDVYCYVDHLDGDVFHITAPSKFTFEEAEAECTSRDARLATVGELQA<br>AWRNGFDQCDYGWLSDASVRHPVTVARAQCGGGLLGVRTLYRFENQTCFPLPDSRFDAYCFGGG<br>GSGGGGSGGGGSEPRVPITQNPCPPLKECPPCAAPDAAGGPSVFIFPPKIKDVLMISLSPMVTCVVV<br>DVSEDDPDVQISWFVNNVEVHTAQTQTHREDYNSTLRVVSALPIQHQDWMSGKEFKCKVNNRALG<br>SPIEKTISKPRGPVRAPQVYVLPPPAEEMTKKEFSLTCMITGFLPAEIAVDWTSNGRTEQNYKNTATV<br>LDSDGSYFMYSKLRVQKSTWERGSLFACSVVHEGLHNHLTTKTISRSLGK |
| Versican S139$G_T$-MSA | TSPPVKGSLSGKVVLPCHFSTLPTLPPNYNTSEFLRIKWSKMEVDKNGKDIKETTVLVAQNGNIKIGQ<br>DYKGRVSVPTHPDDVGDASLTMVKLRASDAAVYRCDVMYGIEDTQDTMSLAVDGVVFHYRAATGR<br>YTLNFAAAQQACLDIGAVIASPEQLFAAYEDGFEQCDAGWLSDQTVRYPIRAPREGCYGDMMGKEG<br>VRTYGFRSPQETYDVYCYVDHLDGDVFHITAPSKFTFEEAEAECTSRDARLATVGELQAAWRNGFD<br>QCDYGWLSDASVRHPVTVARAQCGGGLLGVRTLYRFENQTCFPLPDSRFDAYCFGGGSGGGSEA<br>HKSEIAHRYNDLGEQHFKGLVLIAFSQYLQKCSYDEHAKLVQEVTDFAKTCVADESAANCDKSLHTL<br>FGDKLCAIPNLRENYGELADCCTKQEPERNECFLQHKDDNPSLPPFERPEAEAMCTSFKENPTTFM<br>GHYLHEVARRHPYFYAPELLYYAEQYNEILTQCCAEADKESCLTPKLDGVKEKALVSSVRQRMKCS<br>SMQKFGERAFKAWAVARLSQTFPNADFAEITKLATDLTKVNKECCHGDLLECADDRAELAKYMCEN<br>QATISSKLQTCCDKPLLKKAHCLSEVEHDTMPADLPAIAADFVEDQEVCKNYAEAKDVFLGTFLYEYS<br>RRHPDYSVSLLLRLAKKYEATLEKCCAEANPPACYGTVLAEFQPLVEEPKNLVKTNCDLYEKLGEYG<br>FQNAILVRYTQKAPQVSTPTLVEAARNLGRVGTKCCTLPEDQRLPCVEDYLSAILNRVCLLHEKTPVS<br>EHVTKCCSGSLVERRPCFSALTVDETYVPKEFKAETFTFHSDICTLPEKEKQIKKQTALAELVKHKPK<br>ATAEQLKTVMDDFAQFLDTCCKAADKDTCFSTEGPNLVTRCKDALAHHHHHH |
| Versican S139$G_T$-Fc (mIgG2C LALA-PG) | TSPPVKGSLSGKVVLPCHFSTLPTLPPNYNTSEFLRIKWSKMEVDKNGKDIKETTVLVAQNGNIKIGQ<br>DYKGRVSVPTHPDDVGDASLTMVKLRASDAAVYRCDVMYGIEDTQDTMSLAVDGVVFHYRAATGR<br>YTLNFAAAQQACLDIGAVIASPEQLFAAYEDGFEQCDAGWLSDQTVRYPIRAPREGCYGDMMGKEG<br>VRTYGFRSPQETYDVYCYVDHLDGDVFHITAPSKFTFEEAEAECTSRDARLATVGELQAAWRNGFD<br>QCDYGWLSDASVRHPVTVARAQCGGGLLGVRTLYRFENQTCFPLPDSRFDAYCFGGGGSGGGGS<br>GGGGSEPRVPITQNPCPPLKECPPCAAPDAAGGPSVFIFPPKIKDVLMISLSPMVTCVVVDVSEDDP<br>DVQISWFVNNVEVHTAQTQTHREDYNSTLRVVSALPIQHQDWMSGKEFKCKVNNRALGSPIEKTISK<br>PRGPVRAPQVYVLPPPAEEMTKKEFSLTCMITGFLPAEIAVDWTSNGRTEQNYKNTATVLDSDGSYF<br>MYSKLRVQKSTWERGSLFACSVVHEGLHNHLTTKTISRSLGK |
| Fc-versican (mIgG2C LALA-PG) | EPRVPITQNPCPPLKECPPCAAPDAAGGPSVFIFPPKIKDVLMISLSPMVTCVVVDVSEDDPDVQISW<br>FVNNVEVHTAQTQTHREDYNSTLRVVSALPIQHQDWMSGKEFKCKVNNRALGSPIEKTISKPRGPV<br>RAPQVYVLPPPAEEMTKKEFSLTCMITGFLPAEIAVDWTSNGRTEQNYKNTATVLDSDGSYFMYSKL<br>RVQKSTWERGSLFACSVVHEGLHNHLTTKTISRSLGKGGGGSGGGGSGGGGSLHQAKMETSPPV<br>KGSLSGKVVLPCHFSTLPTLPPNYNTSEFLRIKWSKMEVDKNGKDIKETTVLVAQNGNIKIGQDYKGR<br>VSVPTHPDDVGDASLTMVKLRASDAAVYRCDVMYGIEDTQDTMSLAVDGVVFHYRAATSRYTLNFA<br>AAQQACLDIGAVIASPEQLFAAYEDGFEQCDAGWLSDQTVRYPIRAPREGCYGDMMGKEGVRTYG<br>FRSPQETYDVYCYVDHLDGDVFHITAPSKFTFEEAEAECTSRDARLATVGELQAAWRNGFDQCDYG<br>WLSDASVRHPVTVARAQCGGGLLGVRTLYRFENQTCFPLPDSRFDAYCF |
| Fc-versican S139G (mIgG2C LALA-PG) | EPRVPITQNPCPPLKECPPCAAPDAAGGPSVFIFPPKIKDVLMISLSPMVTCVVVDVSEDDPDVQISW<br>FVNNVEVHTAQTQTHREDYNSTLRVVSALPIQHQDWMSGKEFKCKVNNRALGSPIEKTISKPRGPV<br>RAPQVYVLPPPAEEMTKKEFSLTCMITGFLPAEIAVDWTSNGRTEQNYKNTATVLDSDGSYFMYSKL<br>RVQKSTWERGSLFACSVVHEGLHNHLTTKTISRSLGKGGGGSGGGGSGGGGSLHQAKMETSPPV<br>KGSLSGKVVLPCHFSTLPTLPPNYNTSEFLRIKWSKMEVDKNGKDIKETTVLVAQNGNIKIGQDYKGR<br>VSVPTHPDDVGDASLTMVKLRASDAAVYRCDVMYGIEDTQDTMSLAVDGVVFHYRAATGRYTLNFA<br>AAQQACLDIGAVIASPEQLFAAYEDGFEQCDAGWLSDQTVRYPIRAPREGCYGDMMGKEGVRTYG<br>FRSPQETYDVYCYVDHLDGDVFHITAPSKFTFEEAEAECTSRDARLATVGELQAAWRNGFDQCDYG<br>WLSDASVRHPVTVARAQCGGGLLGVRTLYRFENQTCFPLPDSRFDAYCF |

| | |
|---|---|
| Fc-versican S139G$_T$ (mIgG2C LALA-PG) | EPRVPITQNPCPPLKECPPCAAPDAAGGPSVFIFPPKIKDVLMISLSPMVTCVVVDVSEDDPDVQISW<br>FVNNVEVHTAQTQTHREDYNSTLRVVSALPIQHQDWMSGKEFKCKVNNRALGSPIEKTISKPRGPV<br>RAPQVYVLPPPAEEMTKKEFSLTCMITGFLPAEIAVDWTSNGRTEQNYKNTATVLDSDGSYFMYSKL<br>RVQKSTWERGSLFACSVVHEGLHNHLTTKTISRSLGKGGGGSGGGGSGGGGSTSPPVKGSLSGKV<br>VLPCHFSTLPTLPPNYNTSEFLRIKWSKMEVDKNGKDIKETTVLVAQNGNIKIGQDYKGRVSVPTHPD<br>DVGDASLTMVKLRASDAAVYRCDVMYGIEDTQDTMSLAVDGVVFHYRAATGRYTLNFAAAQQACL<br>DIGAVIASPEQLFAAYEDGFEQCDAGWLSDQTVRYPIRAPREGCYGDMMGKEGVRTYGFRSPQET<br>YDVYCYVDHLDGDVFHITAPSKFTFEEAEAECTSRDARLATVGELQAAWRNGFDQCDYGWLSDAS<br>VRHPVTVARAQCGGGLLGVRTLYRFENQTCFPLPDSRFDAYCF |
| IgG Light Chain (4420; mCκ) | DVVMTQTPLSLPVSLGDQASISCRSSQSLVHSNGNTYLRWYLQKPGQSPKVLIYKVSNRFSGVPDR<br>FSGSGSGTDFTLKISRVEAEDLGVYFCSQSTHVPWTFGGGTKLEIKRADAAPTVSIFPPSSEQLTSG<br>GASVVCFLNNFYPKDINVKWKIDGSERQNGVLNSWTDQDSKDSTYSMSSTLTLTKDEYERHNSYTC<br>EATHKTSTSPIVKSFNRNEC |
| IgG Heavy Chain (4420; mIgG2c LALA-PG) | DVKLDETGGGLVQPGRPMKLSCVASGFTFSDYWMNWVRQSPEKGLEWVAQIRNKPYNYETYYSD<br>SVKGRFTISRDDSKSSVYLQMNNLRVEDMGIYYCTGSYYGMDYWGQGTSVTVSAKTTAPSVYPLAP<br>VCGGTTGSSVTLGCLVKGYFPEPVTLTWNSGSLSSGVHTFPALLQSGLYTLSSSVTVTSNTWPSQTI<br>TCNVAHPASSTKVDKKIEPRVPITQNPCPPLKECPPCAAPDAAGGPSVFIFPPKIKDVLMISLSPMVTC<br>VVVDVSEDDPDVQISWFVNNVEVHTAQTQTHREDYNSTLRVVSALPIQHQDWMSGKEFKCKVNNR<br>ALGSPIEKTISKPRGPVRAPQVYVLPPPAEEMTKKEFSLTCMITGFLPAEIAVDWTSNGRTEQNYKNT<br>ATVLDSDGSYLMYSKLTVQKSTWERGSLFACSVVHEGLHNHLTTKTISRSLGK |
| Lumican-Fc (mIgG2C LALA-PG) | QYYDYDIPLFMYGQISPNCAPECNCPHSYPTAMYCDDLKLKSVPMVPPGIKYLYLRNNQIDHIDEKAF<br>ENVTDLQWLILDHNLLENSKIKGKVFSKLKQLKKLHINYNNLTESVGPLPKSLQDLQLTNNKISKLGSF<br>DGLVNLTFIYLQHNQLKEDAVSASLKGLKSLEYLDLSFNQMSKLPAGLPTSLLTLYLDNNKISNIPDEY<br>FKRFTGLQYLRLSHNELADSGVPGNSFNISSLLELDLSYNKLKSIPTVNENLENYYLEVNELEKFDVKS<br>FCKILGPLSYSKIKHLRLDGNPLTQSSLPPDMYECLRVANEITVNGGGGSGGGGSEPRVPITQNPCP<br>PLKECPPCAAPDAAGGPSVFIFPPKIKDVLMISLSPMVTCVVVDVSEDDPDVQISWFVNNVEVHTAQ<br>TQTHREDYNSTLRVVSALPIQHQDWMSGKEFKCKVNNRALGSPIEKTISKPRGPVRAPQVYVLPPPA<br>EEMTKKEFSLTCMITGFLPAEIAVDWTSNGRTEQNYKNTATVLDSDGSYFMYSKLRVQKSTWERGS<br>LFACSVVHEGLHNHLTTKTISRSLGK |
| IgG-IL15 Heavy Chain (mIgG2C LALA-PG) | DVKLDETGGGLVQPGRPMKLSCVASGFTFSDYWMNWVRQSPEKGLEWVAQIRNKPYNYETYYSD<br>SVKGRFTISRDDSKSSVYLQMNNLRVEDMGIYYCTGSYYGMDYWGQGTSVTVSAKTTAPSVYPLAP<br>VCGGTTGSSVTLGCLVKGYFPEPVTLTWNSGSLSSGVHTFPALLQSGLYTLSSSVTVTSNTWPSQTI<br>TCNVAHPASSTKVDKKIEPRVPITQNPCPPLKECPPCAAPDAAGGPSVFIFPPKIKDVLMISLSPMVTC<br>VVVDVSEDDPDVQISWFVNNVEVHTAQTQTHREDYNSTLRVVSALPIQHQDWMSGKEFKCKVNNR<br>ALGSPIEKTISKPRGPVRAPQVYVLPPPAEEMTKKEFSLTCMITGFLPAEIAVDWTSNGRTEQNYKNT<br>ATVLDSDGSYLMYSKLTVQKSTWERGSLFACSVVHEGLHNHLTTKTISRSLGKGGGGSGGGGSGT<br>TCPPPVSIEHADIRVKNYSVNSRERYVCNSGFKRKAGTSTLIECVINKNTNVAHWTTPSLKCIRDPSL<br>AGGSGGSGGSGGSGGSGGSGGNWIDVRYDLEKIESLIQSIHIDTTLYTDSDFHPSCKVTAMNCFLLE<br>LQVILHEYSNMTLNETVRNVLYLANSTLSSNKNVAESGCKECEELEEKTFTEFLQSFIRIVQMFINTS |
| IL12-IgG Heavy Chain (mIgG2C LALA-PG) | MWELEKDVYVVEVDWTPDAPGETVNLTCDTPEEDDITWTSDQRHGVIGSGKTLTITVKEFLDAGQY<br>TCHKGGETLSHSHLLLHKKENGIWSTEILKNFKNKTFLKCEAPNYSGRFTCSWLVQRNMDLKFNIKS<br>SSSSPDSRAVTCGMASLSAEKVTLDQRDYEKYSVSCQEDVTCPTAEETLPIELALEARQQNKYENY<br>STSFFIRDIIKPDPPKNLQMKPLKNSQVEVSWEYPDSWSTPHSYFSLKFFVRIQRKKEKMKETEEGC<br>NQKGAFLVEKTSTEVQCKGGNVCVQAQDRYYNSSCSKWACVPCRVRSGGSGGGSGGGSGGGSR<br>VIPVSGPARCLSQSRNLLKTTDDMVKTAREKLKHYSCTAEDIDHEDITRDQTSTLKTCLPLELHKNES<br>CLATRETSSTTRGSCLPPQKTSLMMTLCLGSIYEDLKMYQTEFQAINAALQNHNHQQIILDKGMLVAI<br>DELMQSLNHNGETLRQKPPVGEADPYRVKMKLCILLHAFSTRVVTINRVMGYLSSAGGGGSGGGG<br>SGGGGSDVKLDETGGGLVQPGRPMKLSCVASGFTFSDYWMNWVRQSPEKGLEWVAQIRNKPYNY<br>ETYYSDSVKGRFTISRDDSKSSVYLQMNNLRVEDMGIYYCTGSYYGMDYWGQGTSVTVSAKTTAP<br>SVYPLAPVCGGTTGSSVTLGCLVKGYFPEPVTLTWNSGSLSSGVHTFPALLQSGLYTLSSSVTVTSN<br>TWPSQTITCNVAHPASSTKVDKKIEPRVPITQNPCPPLKECPPCAAPDAAGGPSVFIFPPKIKDVLMIS<br>LSPMVTCVVVDVSEDDPDVQISWFVNNVEVHTAQTQTHREDYNSTLRVVSALPIQHQDWMSGKEF<br>KCKVNNRALGSPIEKTISKPRGPVRAPQVYVLPPPAEEMTKKEFSLTCMITGFLPAEIAVDWTSNGRT<br>EQNYKNTATVLDSDGSYLMYSKLTVQKSTWERGSLFACSVVHEGLHNHLTTKTISRSLGK |

| | |
|---|---|
| VHH$_2$-MSA-IL15 | QVQLVETGGGLVQPGESLRLSCVASGFTLDHSAVGWFRQVPGKEREKLLCINANGVSLDYADSIKG<br>RFTISRDNAKNTVYLQMNDLKPEDTATYSCAATREFCSAYVFLYEHWGQGTQVTVSSGGGSGGGS<br>GGGSQVQLVETGGGLVQPGESLRLSCVASGFTLDHSAVGWFRQVPGKEREKLLCINANGVSLDYA<br>DSIKGRFTISRDNAKNTVYLQMNDLKPEDTATYSCAATREFCSAYVFLYEHWGQGTQVTVSSGGGS<br>GGGSGGGSEAHKSEIAHRYNDLGEQHFKGLVLIAFSQYLQKCSYDEHAKLVQEVTDFAKTCVADES<br>AANCDKSLHTLFGDKLCAIPNLRENYGELADCCTKQEPERNECFLQHKDDNPSLPPFERPEAEAMC<br>TSFKENPTTFMGHYLHEVARRHPYFYAPELLYYAEQYNEILTQCCAEADKESCLTPKLDGVKEKALV<br>SSVRQRMKCSSMQKFGERAFKAWAVARLSQTFPNADFAEITKLATDLTKVNKECCHGDLLECADDR<br>AELAKYMCENQATISSKLQTCCDKPLLKKAHCLSEVEHDTMPADLPAIAADFVEDQEVCKNYAEAKD<br>VFLGTFLYEYSRRHPDYSVSLLLRLAKKYEATLEKCCAEANPPACYGTVLAEFQPLVEEPKNLVKTN<br>CDLYEKLGEYGFQNAILVRYTQKAPQVSTPTLVEAARNLGRVGTKCCTLPEDQRLPCVEDYLSAILN<br>RVCLLHEKTPVSEHVTKCCSGSLVERRPCFSALTVDETYVPKEFKAETFTFHSDICTLPEKEKQIKKQ<br>TALAELVKHKPKATAEQLKTVMDDFAQFLDTCCKAADKDTCFSTEGPNLVTRCKDALAGGGSGTTC<br>PPPVSIEHADIRVKNYSVNSRERYVCNSGFKRKAGTSTLIECVINKNTNVAHWTTPSLKCIRDPSLAG<br>GSGGSGGSGGSGGSGGSGGNWIDVRYDLEKIESLIQSIHIDTTLYTDSDFHPSCKVTAMNCFLLELQ<br>VILHEYSNMTLNETVRNVLYLANSTLSSNKNVAESGCKECEELEEKTFTEFLQSFIRIVQMFINTSHHH<br>HHH |
| IL12-VHH$_2$-MSA | MWELEKDVYVVEVDWTPDAPGETVNLTCDTPEEDDITWTSDQRHGVIGSGKTLTITVKEFLDAGQY<br>TCHKGGETLSHSHLLLHKKENGIWSTEILKNFKNKTFLKCEAPNYSGRFTCSWLVQRNMDLKFNIKS<br>SSSSPDSRAVTCGMASLSAEKVTLDQRDYEKYSVSCQEDVTCPTAEETLPIELALEARQQNKYENY<br>STSFFIRDIIKPDPPKNLQMKPLKNSQVEVSWEYPDSWSTPHSYFSLKFFVRIQRKKEKMKETEEGC<br>NQKGAFLVEKTSTEVQCKGGNVCVQAQDRYYNSSCSKWACVPCRVRSGGSGGGSGGGSGGGSR<br>VIPVSGPARCLSQSRNLLKTTDDMVKTAREKLKHYSCTAEDIDHEDITRDQTSTLKTCLPLELHKNES<br>CLATRETSSTTRGSCLPPQKTSLMMTLCLGSIYEDLKMYQTEFQAINAALQNHNHQQIILDKGMLVAI<br>DELMQSLNHNGETLRQKPPVGEADPYRVKMKLCILLHAFSTRVVTINRVMGYLSSAGGGSGGGSG<br>GGSQVQLVETGGGLVQPGESLRLSCVASGFTLDHSAVGWFRQVPGKEREKLLCINANGVSLDYAD<br>SIKGRFTISRDNAKNTVYLQMNDLKPEDTATYSCAATREFCSAYVFLYEHWGQGTQVTVSSGGGSG<br>GGSGGGSQVQLVETGGGLVQPGESLRLSCVASGFTLDHSAVGWFRQVPGKEREKLLCINANGVSL<br>DYADSIKGRFTISRDNAKNTVYLQMNDLKPEDTATYSCAATREFCSAYVFLYEHWGQGTQVTVSSG<br>GGSGGGSGGGSEAHKSEIAHRYNDLGEQHFKGLVLIAFSQYLQKCSYDEHAKLVQEVTDFAKTCVA<br>DESAANCDKSLHTLFGDKLCAIPNLRENYGELADCCTKQEPERNECFLQHKDDNPSLPPFERPEAE<br>AMCTSFKENPTTFMGHYLHEVARRHPYFYAPELLYYAEQYNEILTQCCAEADKESCLTPKLDGVKEK<br>ALVSSVRQRMKCSSMQKFGERAFKAWAVARLSQTFPNADFAEITKLATDLTKVNKECCHGDLLECA<br>DDRAELAKYMCENQATISSKLQTCCDKPLLKKAHCLSEVEHDTMPADLPAIAADFVEDQEVCKNYAE<br>AKDVFLGTFLYEYSRRHPDYSVSLLLRLAKKYEATLEKCCAEANPPACYGTVLAEFQPLVEEPKNLV<br>KTNCDLYEKLGEYGFQNAILVRYTQKAPQVSTPTLVEAARNLGRVGTKCCTLPEDQRLPCVEDYLSA<br>ILNRVCLLHEKTPVSEHVTKCCSGSLVERRPCFSALTVDETYVPKEFKAETFTFHSDICTLPEKEKQIK<br>KQTALAELVKHKPKATAEQLKTVMDDFAQFLDTCCKAADKDTCFSTEGPNLVTRCKDALAHHHHHH |
| Lumican-MSA-IL15 | QYYDYDIPLFMYGQISPNCAPECNCPHSYPTAMYCDDLKLKSVPMVPPGIKYLYLRNNQIDHIDEKAF<br>ENVTDLQWLILDHNLLENSKIKGKVFSKLKQLKKLHINYNNLTESVGPLPKSLQDLQLTNNKISKLGSF<br>DGLVNLTFIYLQHNQLKEDAVSASLKGLKSLEYLDLSFNQMSKLPAGLPTSLLTLYLDNNKISNIPDEY<br>FKRFTGLQYLRLSHNELADSGVPGNSFNISSLLELDLSYNKLKSIPTVNENLENYYLEVNELEKFDVKS<br>FCKILGPLSYSKIKHLRLDGNPLTQSSLPPDMYECLRVANEITVNGGGSGGGSEAHKSEIAHRYNDL<br>GEQHFKGLVLIAFSQYLQKCSYDEHAKLVQEVTDFAKTCVADESAANCDKSLHTLFGDKLCAIPNLR<br>ENYGELADCCTKQEPERNECFLQHKDDNPSLPPFERPEAEAMCTSFKENPTTFMGHYLHEVARRH<br>PYFYAPELLYYAEQYNEILTQCCAEADKESCLTPKLDGVKEKALVSSVRQRMKCSSMQKFGERAFK<br>AWAVARLSQTFPNADFAEITKLATDLTKVNKECCHGDLLECADDRAELAKYMCENQATISSKLQTCC<br>DKPLLKKAHCLSEVEHDTMPADLPAIAADFVEDQEVCKNYAEAKDVFLGTFLYEYSRRHPDYSVSLL<br>LRLAKKYEATLEKCCAEANPPACYGTVLAEFQPLVEEPKNLVKTNCDLYEKLGEYGFQNAILVRYTQ<br>KAPQVSTPTLVEAARNLGRVGTKCCTLPEDQRLPCVEDYLSAILNRVCLLHEKTPVSEHVTKCCSGS<br>LVERRPCFSALTVDETYVPKEFKAETFTFHSDICTLPEKEKQIKKQTALAELVKHKPKATAEQLKTVM<br>DDFAQFLDTCCKAADKDTCFSTEGPNLVTRCKDALAGGGSGGGSGTTCPPPVSIEHADIRVKNYSV<br>NSRERYVCNSGFKRKAGTSTLIECVINKNTNVAHWTTPSLKCIRDPSLAGGSGGSGGSGGSGGSGG<br>SGGNWIDVRYDLEKIESLIQSIHIDTTLYTDSDFHPSCKVTAMNCFLLELQVILHEYSNMTLNETVRNV<br>LYLANSTLSSNKNVAESGCKECEELEEKTFTEFLQSFIRIVQMFINTSHHHHHH |
| IL12-lumican-MSA | MWELEKDVYVVEVDWTPDAPGETVNLTCDTPEEDDITWTSDQRHGVIGSGKTLTITVKEFLDAGQY<br>TCHKGGETLSHSHLLLHKKENGIWSTEILKNFKNKTFLKCEAPNYSGRFTCSWLVQRNMDLKFNIKS<br>SSSSPDSRAVTCGMASLSAEKVTLDQRDYEKYSVSCQEDVTCPTAEETLPIELALEARQQNKYENY |

| | |
|---|---|
| | STSFFIRDIIKPDPPKNLQMKPLKNSQVEVSWEYPDSWSTPHSYFSLKFFVRIQRKKEKMKETEEGC<br>NQKGAFLVEKTSTEVQCKGGNVCVQAQDRYYNSSCSKWACVPCRVRSGGSGGGSGGGSGGGSR<br>VIPVSGPARCLSQSRNLLKTTDDMVKTAREKLKHYSCTAEDIDHEDITRDQTSTLKTCLPLELHKNES<br>CLATRETSSTTRGSCLPPQKTSLMMTLCLGSIYEDLKMYQTEFQAINAALQNHNHQQIILDKGMLVAI<br>DELMQSLNHNGETLRQKPPVGEADPYRVKMKLCILLHAFSTRVVTINRVMGYLSSAGGGSGGGSG<br>GGSQYYDYDIPLFMYGQISPNCAPECNCPHSYPTAMYCDDLKLKSVPMVPPGIKYLYLRNNQIDHID<br>EKAFENVTDLQWLILDHNLLENSKIKGKVFSKLKQLKKLHINYNNLTESVGPLPKSLQDLQLTNNKISK<br>LGSFDGLVNLTFIYLQHNQLKEDAVSASLKGLKSLEYLDLSFNQMSKLPAGLPTSLLTLYLDNNKISNI<br>PDEYFKRFTGLQYLRLSHNELADSGVPGNSFNISSLLELDLSYNKLKSIPTVNENLENYYLEVNELEKF<br>DVKSFCKILGPLSYSKIKHLRLDGNPLTQSSLPPDMYECLRVANEITVNGGGSGGGSEAHKSEIAHRY<br>NDLGEQHFKGLVLIAFSQYLQKCSYDEHAKLVQEVTDFAKTCVADESAANCDKSLHTLFGDKLCAIP<br>NLRENYGELADCCTKQEPERNECFLQHKDDNPSLPPFERPEAEAMCTSFKENPTTFMGHYLHEVA<br>RRHPYFYAPELLYYAEQYNEILTQCCAEADKESCLTPKLDGVKEKALVSSVRQRMKCSSMQKFGER<br>AFKAWAVARLSQTFPNADFAEITKLATDLTKVNKECCHGDLLECADDRAELAKYMCENQATISSKLQ<br>TCCDKPLLKKAHCLSEVEHDTMPADLPAIAADFVEDQEVCKNYAEAKDVFLGTFLYEYSRRHPDYSV<br>SLLLRLAKKYEATLEKCCAEANPPACYGTVLAEFQPLVEEPKNLVKTNCDLYEKLGEYGFQNAILVRY<br>TQKAPQVSTPTLVEAARNLGRVGTKCCTLPEDQRLPCVEDYLSAILNRVCLLHEKTPVSEHVTKCCS<br>GSLVERRPCFSALTVDETYVPKEFKAETFTFHSDICTLPEKEKQIKKQTALAELVKHKPKATAEQLKTV<br>MDDFAQFLDTCCKAADKDTCFSTEGPNLVTRCKDALAHHHHHH |
| Lumican-Fc-IL15 (mIgG2C LALA-PG) | QYYDYDIPLFMYGQISPNCAPECNCPHSYPTAMYCDDLKLKSVPMVPPGIKYLYLRNNQIDHIDEKAF<br>ENVTDLQWLILDHNLLENSKIKGKVFSKLKQLKKLHINYNNLTESVGPLPKSLQDLQLTNNKISKLGSF<br>DGLVNLTFIYLQHNQLKEDAVSASLKGLKSLEYLDLSFNQMSKLPAGLPTSLLTLYLDNNKISNIPDEY<br>FKRFTGLQYLRLSHNELADSGVPGNSFNISSLLELDLSYNKLKSIPTVNENLENYYLEVNELEKFDVKS<br>FCKILGPLSYSKIKHLRLDGNPLTQSSLPPDMYECLRVANEITVNGGGGSGGGGSEPRVPITQNPCP<br>PLKECPPCAAPDAAGGPSVFIFPPKIKDVLMISLSPMVTCVVVDVSEDDPDVQISWFVNNVEVHTAQ<br>TQTHREDYNSTLRVVSALPIQHQDWMSGKEFKCKVNNRALGSPIEKTISKPRGPVRAPQVYVLPPPA<br>EEMTKKEFSLTCMITGFLPAEIAVDWTSNGRTEQNYKNTATVLDSDGSYFMYSKLRVQKSTWERGS<br>LFACSVVHEGLHNHLTTKTISRSLGKGGGSGGGSGTTCPPPVSIEHADIRVKNYSVNSRERYVCNSG<br>FKRKAGTSTLIECVINKNTNVAHWTTPSLKCIRDPSLAGGSGGSGGSGGSGGSGGSGGNWIDVRYD<br>LEKIESLIQSIHIDTTLYTDSDFHPSCKVTAMNCFLLELQVILHEYSNMTLNETVRNVLYLANSTLSSNK<br>NVAESGCKECEELEEKTFTEFLQSFIRIVQMFINTS |
| IL12-lumican-Fc (mIgG2C LALA-PG) | MWELEKDVYVVEVDWTPDAPGETVNLTCDTPEEDDITWTSDQRHGVIGSGKTLTITVKEFLDAGQY<br>TCHKGGETLSHSHLLLHKKENGIWSTEILKNFKNKTFLKCEAPNYSGRFTCSWLVQRNMDLKFNIKS<br>SSSSPDSRAVTCGMASLSAEKVTLDQRDYEKYSVSCQEDVTCPTAEETLPIELALEARQQNKYENY<br>STSFFIRDIIKPDPPKNLQMKPLKNSQVEVSWEYPDSWSTPHSYFSLKFFVRIQRKKEKMKETEEGC<br>NQKGAFLVEKTSTEVQCKGGNVCVQAQDRYYNSSCSKWACVPCRVRSGGSGGGSGGGSGGGSR<br>VIPVSGPARCLSQSRNLLKTTDDMVKTAREKLKHYSCTAEDIDHEDITRDQTSTLKTCLPLELHKNES<br>CLATRETSSTTRGSCLPPQKTSLMMTLCLGSIYEDLKMYQTEFQAINAALQNHNHQQIILDKGMLVAI<br>DELMQSLNHNGETLRQKPPVGEADPYRVKMKLCILLHAFSTRVVTINRVMGYLSSAGGGSGGGSG<br>GGSQYYDYDIPLFMYGQISPNCAPECNCPHSYPTAMYCDDLKLKSVPMVPPGIKYLYLRNNQIDHID<br>EKAFENVTDLQWLILDHNLLENSKIKGKVFSKLKQLKKLHINYNNLTESVGPLPKSLQDLQLTNNKISK<br>LGSFDGLVNLTFIYLQHNQLKEDAVSASLKGLKSLEYLDLSFNQMSKLPAGLPTSLLTLYLDNNKISNI<br>PDEYFKRFTGLQYLRLSHNELADSGVPGNSFNISSLLELDLSYNKLKSIPTVNENLENYYLEVNELEKF<br>DVKSFCKILGPLSYSKIKHLRLDGNPLTQSSLPPDMYECLRVANEITVNGGGGSGGGGSGGGGSEP<br>RVPITQNPCPPLKECPPCAAPDAAGGPSVFIFPPKIKDVLMISLSPMVTCVVVDVSEDDPDVQISWFV<br>NNVEVHTAQTQTHREDYNSTLRVVSALPIQHQDWMSGKEFKCKVNNRALGSPIEKTISKPRGPVRA<br>PQVYVLPPPAEEMTKKEFSLTCMITGFLPAEIAVDWTSNGRTEQNYKNTATVLDSDGSYFMYSKLRV<br>QKSTWERGSLFACSVVHEGLHNHLTTKTISRSLGK |
| Versican $S139G_T$-MSA-IL15 | TSPPVKGSLSGKVVLPCHFSTLPTLPPNYNTSEFLRIKWSKMEVDKNGKDIKETTVLVAQNGNIKIGQ<br>DYKGRVSVPTHPDDVGDASLTMVKLRASDAAVYRCDVMYGIEDTQDTMSLAVDGVVFHYRAATGR<br>YTLNFAAAQQACLDIGAVIASPEQLFAAYEDGFEQCDAGWLSDQTVRYPIRAPREGCYGDMMGKEG<br>VRTYGFRSPQETYDVYCYVDHLDGDVFHITAPSKFTFEEAEAECTSRDARLATVGELQAAWRNGFD<br>QCDYGWLSDASVRHPVTVARAQCGGGLLGVRTLYRFENQTCFPLPDSRFDAYCFGGGSGGGSEA<br>HKSEIAHRYNDLGEQHFKGLVLIAFSQYLQKCSYDEHAKLVQEVTDFAKTCVADESAANCDKSLHTL<br>FGDKLCAIPNLRENYGELADCCTKQEPERNECFLQHKDDNPSLPPFERPEAEAMCTSFKENPTTFM<br>GHYLHEVARRHPYFYAPELLYYAEQYNEILTQCCAEADKESCLTPKLDGVKEKALVSSVRQRMKCS<br>SMQKFGERAFKAWAVARLSQTFPNADFAEITKLATDLTKVNKECCHGDLLECADDRAELAKYMCEN<br>QATISSKLQTCCDKPLLKKAHCLSEVEHDTMPADLPAIAADFVEDQEVCKNYAEAKDVFLGTFLYEYS |

| | |
|---|---|
| | RRHPDYSVSLLLRLAKKYEATLEKCCAEANPPACYGTVLAEFQPLVEEPKNLVKTNCDLYEKLGEYG<br>FQNAILVRYTQKAPQVSTPTLVEAARNLGRVGTKCCTLPEDQRLPCVEDYLSAILNRVCLLHEKTPVS<br>EHVTKCCSGSLVERRPCFSALTVDETYVPKEFKAETFTFHSDICTLPEKEKQIKKQTALAELVKHKPK<br>ATAEQLKTVMDDFAQFLDTCCKAADKDTCFSTEGPNLVTRCKDALAGGGSGTTCPPPVSIEHADIRV<br>KNYSVNSRERYVCNSGFKRKAGTSTLIECVINKNTNVAHWTTPSLKCIRDPSLAGGSGGSGGSGGS<br>GGSGGSGGNWIDVRYDLEKIESLIQSIHIDTTLYTDSDFHPSCKVTAMNCFLLELQVILHEYSNMTLNE<br>TVRNVLYLANSTLSSNKNVAESGCKECEELEEKTFTEFLQSFIRIVQMFINTSHHHHHH |
| IL12-versican $S139G_T$-MSA | MWELEKDVYVVEVDWTPDAPGETVNLTCDTPEEDDITWTSDQRHGVIGSGKTLTITVKEFLDAGQY<br>TCHKGGETLSHSHLLLHKKENGIWSTEILKNFKNKTFLKCEAPNYSGRFTCSWLVQRNMDLKFNIKS<br>SSSSPDSRAVTCGMASLSAEKVTLDQRDYEKYSVSCQEDVTCPTAEETLPIELALEARQQNKYENY<br>STSFFIRDIIKPDPPKNLQMKPLKNSQVEVSWEYPDSWSTPHSYFSLKFFVRIQRKKEKMKETEEGC<br>NQKGAFLVEKTSTEVQCKGGNVCVQAQDRYYNSSCSKWACVPCRVRSGGSGGGSGGGSGGGSR<br>VIPVSGPARCLSQSRNLLKTTDDMVKTAREKLKHYSCTAEDIDHEDITRDQTSTLKTCLPLELHKNES<br>CLATRETSSTTRGSCLPPQKTSLMMTLCLGSIYEDLKMYQTEFQAINAALQNHNHQQIILDKGMLVAI<br>DELMQSLNHNGETLRQKPPVGEADPYRVKMKLCILLHAFSTRVVTINRVMGYLSSAGGGSGGGSG<br>GGSTSPPVKGSLSGKVVLPCHFSTLPTLPPNYNTSEFLRIKWSKMEVDKNGKDIKETTVLVAQNGNI<br>KIGQDYKGRVSVPTHPDDVGDASLTMVKLRASDAAVYRCDVMYGIEDTQDTMSLAVDGVVFHYRAA<br>TGRYTLNFAAAQQACLDIGAVIASPEQLFAAYEDGFEQCDAGWLSDQTVRYPIRAPREGCYGDMMG<br>KEGVRTYGFRSPQETYDVYCYVDHLDGDVFHITAPSKFTFEEAEAECTSRDARLATVGELQAAWRN<br>GFDQCDYGWLSDASVRHPVTVARAQCGGGLLGVRTLYRFENQTCFPLPDSRFDAYCFGGGSGGG<br>SEAHKSEIAHRYNDLGEQHFKGLVLIAFSQYLQKCSYDEHAKLVQEVTDFAKTCVADESAANCDKSL<br>HTLFGDKLCAIPNLRENYGELADCCTKQEPERNECFLQHKDDNPSLPPFERPEAEAMCTSFKENPTT<br>FMGHYLHEVARRHPYFYAPELLYYAEQYNEILTQCCAEADKESCLTPKLDGVKEKALVSSVRQRMK<br>CSSMQKFGERAFKAWAVARLSQTFPNADFAEITKLATDLTKVNKECCHGDLLECADDRAELAKYMC<br>ENQATISSKLQTCCDKPLLKKAHCLSEVEHDTMPADLPAIAADFVEDQEVCKNYAEAKDVFLGTFLYE<br>YSRRHPDYSVSLLLRLAKKYEATLEKCCAEANPPACYGTVLAEFQPLVEEPKNLVKTNCDLYEKLGE<br>YGFQNAILVRYTQKAPQVSTPTLVEAARNLGRVGTKCCTLPEDQRLPCVEDYLSAILNRVCLLHEKTP<br>VSEHVTKCCSGSLVERRPCFSALTVDETYVPKEFKAETFTFHSDICTLPEKEKQIKKQTALAELVKHK<br>PKATAEQLKTVMDDFAQFLDTCCKAADKDTCFSTEGPNLVTRCKDALAHHHHHH |
| Versican $S139G_T$-Fc-IL15 | TSPPVKGSLSGKVVLPCHFSTLPTLPPNYNTSEFLRIKWSKMEVDKNGKDIKETTVLVAQNGNIKIGQ<br>DYKGRVSVPTHPDDVGDASLTMVKLRASDAAVYRCDVMYGIEDTQDTMSLAVDGVVFHYRAATSR<br>YTLNFAAAQQACLDIGAVIASPEQLFAAYEDGFEQCDAGWLSDQTVRYPIRAPREGCYGDMMGKEG<br>VRTYGFRSPQETYDVYCYVDHLDGDVFHITAPSKFTFEEAEAECTSRDARLATVGELQAAWRNGFD<br>QCDYGWLSDASVRHPVTVARAQCGGGLLGVRTLYRFENQTCFPLPDSRFDAYCFGGGGSEPRVPI<br>TQNPCPPLKECPPCAAPDAAGGPSVFIFPPKIKDVLMISLSPMVTCVVVDVSEDDPDVQISWFVNNV<br>EVHTAQTQTHREDYNSTLRVVSALPIQHQDWMSGKEFKCKVNNRALGSPIEKTISKPRGPVRAPQV<br>YVLPPPAEEMTKKEFSLTCMITGFLPAEIAVDWTSNGRTEQNYKNTATVLDSDGSYFMYSKLRVQKS<br>TWERGSLFACSVVHEGLHNHLTTKTISRSLGKGGGGSGTTCPPPVSIEHADIRVKNYSVNSRERYVC<br>NSGFKRKAGTSTLIECVINKNTNVAHWTTPSLKCIRDPSLAGGSGGSGGSGGSGGSGGSGGNWIDV<br>RYDLEKIESLIQSIHIDTTLYTDSDFHPSCKVTAMNCFLLELQVILHEYSNMTLNETVRNVLYLANSTLS<br>SNKNVAESGCKECEELEEKTFTEFLQSFIRIVQMFINTS |
| IL12-versican $S139G_T$-Fc | MWELEKDVYVVEVDWTPDAPGETVNLTCDTPEEDDITWTSDQRHGVIGSGKTLTITVKEFLDAGQY<br>TCHKGGETLSHSHLLLHKKENGIWSTEILKNFKNKTFLKCEAPNYSGRFTCSWLVQRNMDLKFNIKS<br>SSSSPDSRAVTCGMASLSAEKVTLDQRDYEKYSVSCQEDVTCPTAEETLPIELALEARQQNKYENY<br>STSFFIRDIIKPDPPKNLQMKPLKNSQVEVSWEYPDSWSTPHSYFSLKFFVRIQRKKEKMKETEEGC<br>NQKGAFLVEKTSTEVQCKGGNVCVQAQDRYYNSSCSKWACVPCRVRSGGSGGGSGGGSGGGSR<br>VIPVSGPARCLSQSRNLLKTTDDMVKTAREKLKHYSCTAEDIDHEDITRDQTSTLKTCLPLELHKNES<br>CLATRETSSTTRGSCLPPQKTSLMMTLCLGSIYEDLKMYQTEFQAINAALQNHNHQQIILDKGMLVAI<br>DELMQSLNHNGETLRQKPPVGEADPYRVKMKLCILLHAFSTRVVTINRVMGYLSSAGGGSGGGSG<br>GGSTSPPVKGSLSGKVVLPCHFSTLPTLPPNYNTSEFLRIKWSKMEVDKNGKDIKETTVLVAQNGNI<br>KIGQDYKGRVSVPTHPDDVGDASLTMVKLRASDAAVYRCDVMYGIEDTQDTMSLAVDGVVFHYRAA<br>TGRYTLNFAAAQQACLDIGAVIASPEQLFAAYEDGFEQCDAGWLSDQTVRYPIRAPREGCYGDMMG<br>KEGVRTYGFRSPQETYDVYCYVDHLDGDVFHITAPSKFTFEEAEAECTSRDARLATVGELQAAWRN<br>GFDQCDYGWLSDASVRHPVTVARAQCGGGLLGVRTLYRFENQTCFPLPDSRFDAYCFGGGGSGG<br>GGSGGGGSEPRVPITQNPCPPLKECPPCAAPDAAGGPSVFIFPPKIKDVLMISLSPMVTCVVVDVSE<br>DDPDVQISWFVNNVEVHTAQTQTHREDYNSTLRVVSALPIQHQDWMSGKEFKCKVNNRALGSPIEK<br>TISKPRGPVRAPQVYVLPPPAEEMTKKEFSLTCMITGFLPAEIAVDWTSNGRTEQNYKNTATVLDSD<br>GSYFMYSKLRVQKSTWERGSLFACSVVHEGLHNHLTTKTISRSLGK |

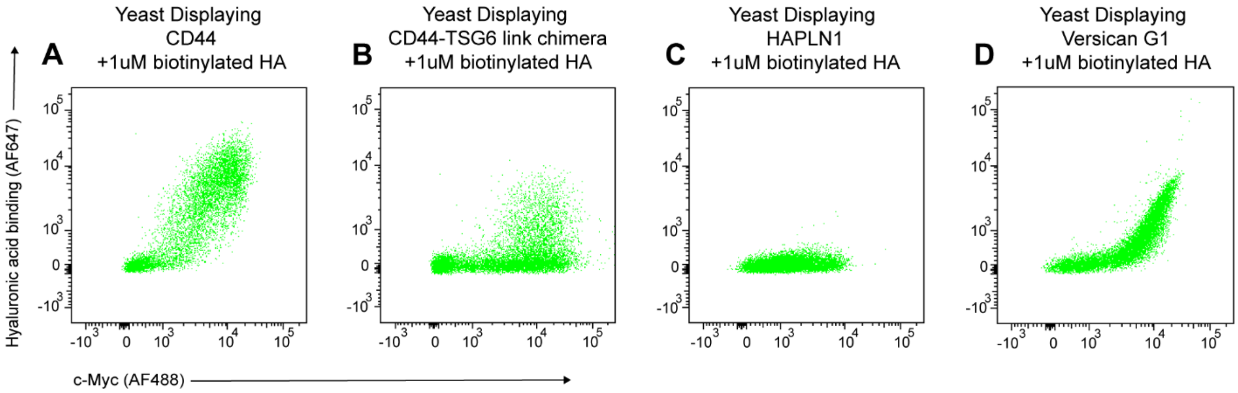


**Supplementary Figure S1**

Versican G1 demonstrates stable expression and hyaluronic acid binding on the yeast surface. **A-D,** Flow cytometry plots of yeast surface displayed (**A**) CD44, (**B**) CD44-TSG6 link chimera, (**C**) HAPLN1, and (**D**) versican G1 stained for display via c-Myc and hyaluronic acid binding.

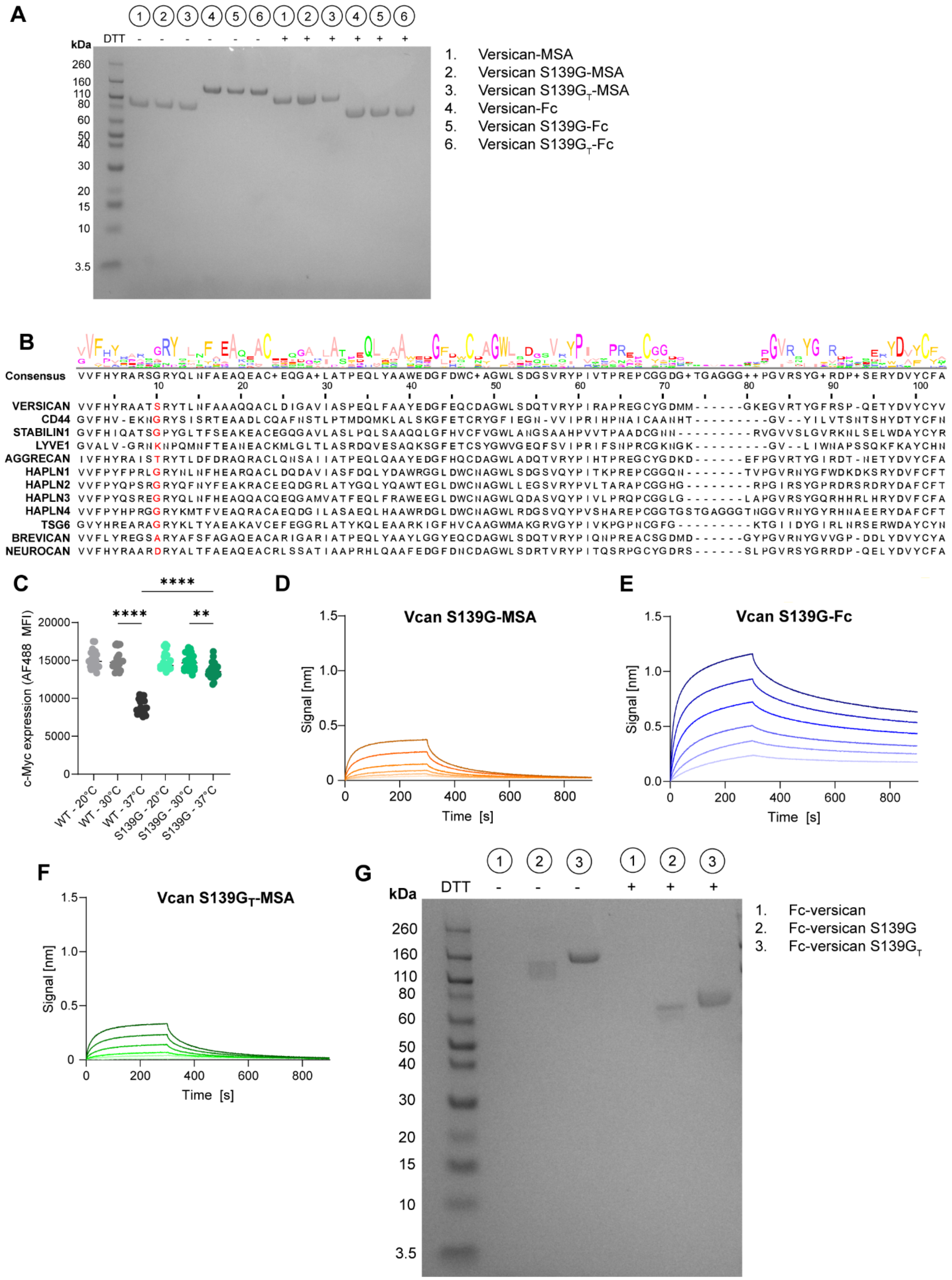


**Supplementary Figure S2**

S139G mutation and N-terminal truncation enhances expression, thermal stability, and binding. **A**, Versican fusions were expressed in the Expi293 express system. Protein A or Nickel-NTA purified fractions are shown on SDS-PAGE. **B**, Multiple sequence alignment of the first Link module in select HA binding proteins. Protein sequences were aligned using Clustal Omega and visualized in JalView. **C**, Clonal yeast expressing wildtype or mutant versican were induced at 20°C, 30°C, or 37°C. Full-length surface expression of protein of interest was assessed with anti-c-Myc antibody (AF488). **D-F,** BLI demonstrating binding kinetics of (**D**) versican S139G-MSA, (**E**) versican S139G-Fc, (**F**) and versican $S139G_{T}$-MSA. (**G**) SDS-PAGE of N-terminal Fc fusions of versican, demonstrating poor expression of fusions without the N-terminal truncation. Statistics: one-way ANOVA followed by Tukey’s multiple-comparison test. **P < 0.01; ****P < 0.0001.

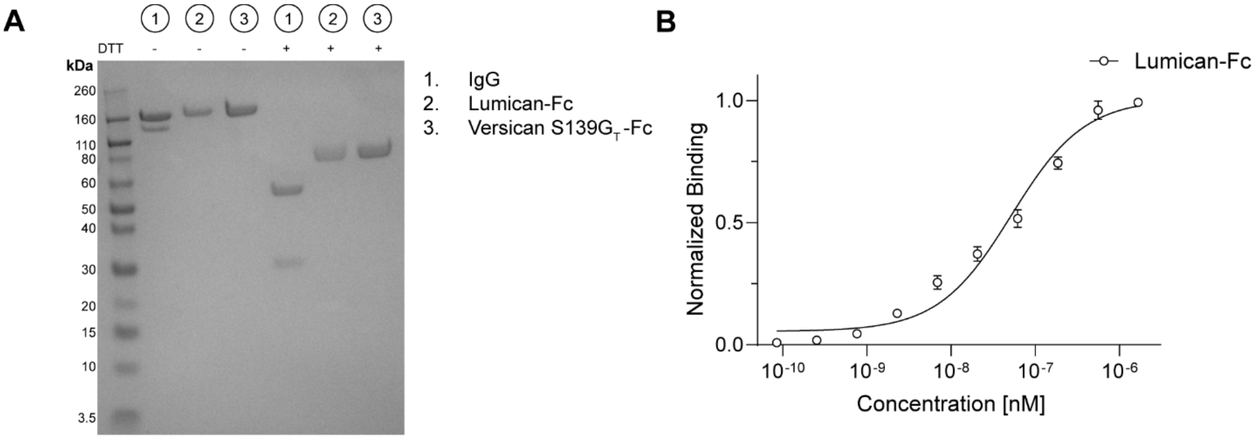


**Supplementary Figure S3**

Characterization of biodistribution proteins. **A**, SDS-PAGE of Protein A purified proteins used for biodistribution work in Figure 2. **B**, Collagen I ELISA of Lumican-Fc shows maintenance of collagen binding in dimer construct.

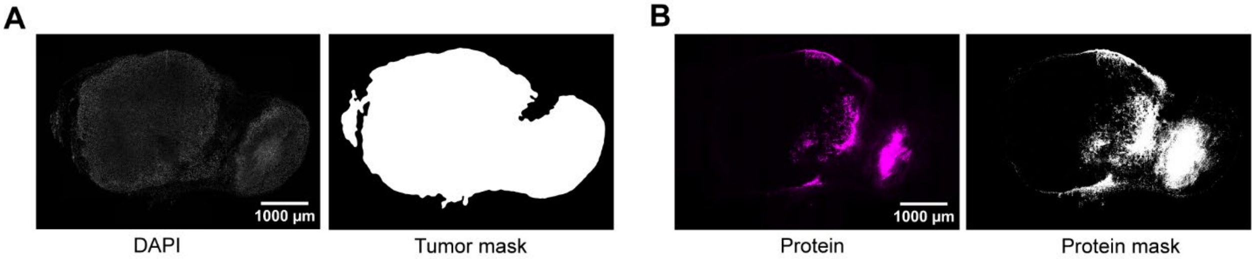


**Supplementary Figure S4**

Analysis pipeline of immunofluorescent microscopy. **A**, Raw DAPI images (left) were used to demarcate tumor boundaries to create a tumor mask (right). **B**, A threshold based on background AF647 signal from uninjected tumors was used to create binary protein signal masks (right) from raw AF647 images (left). All image analysis was performed in ImageJ.

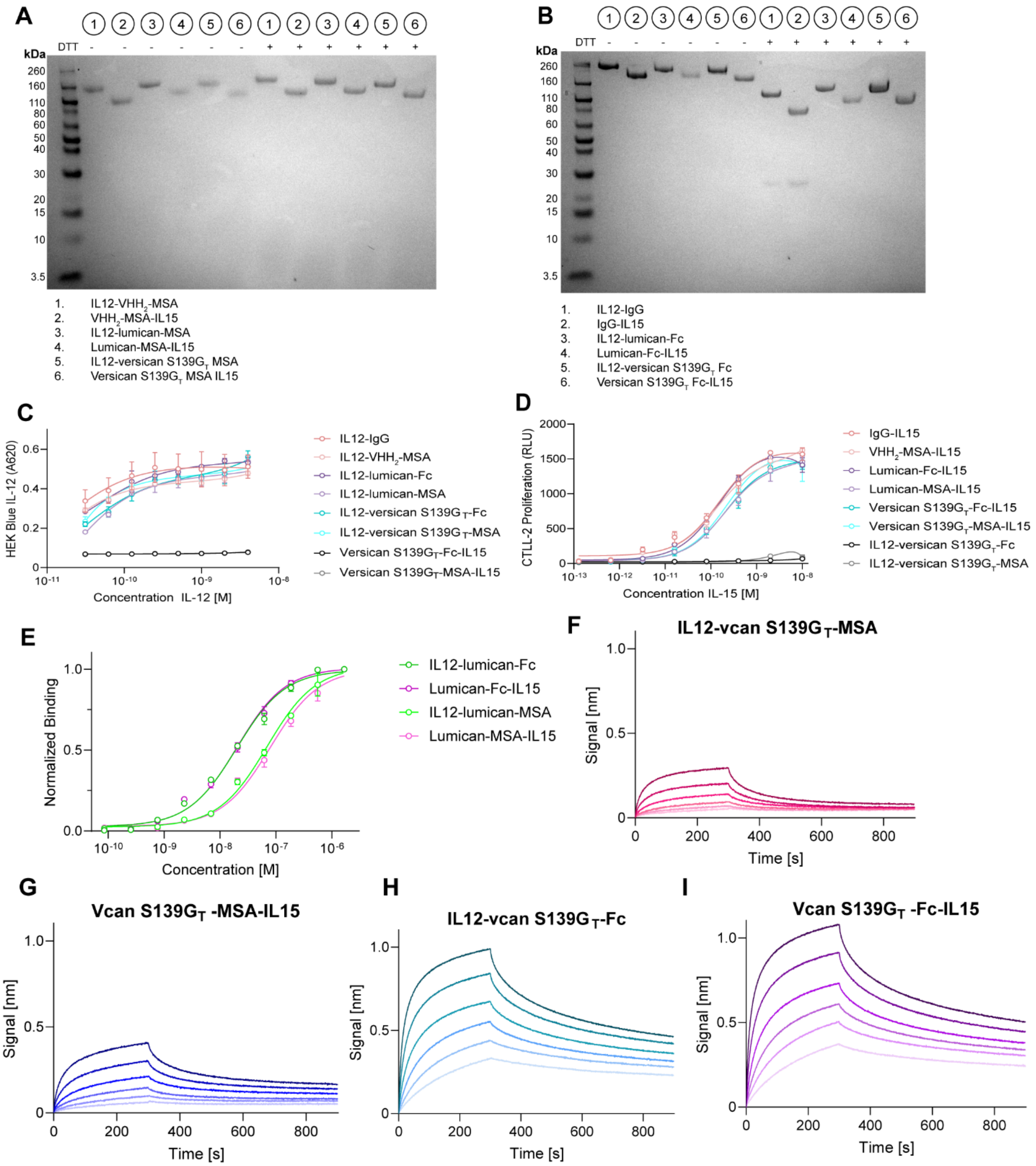


**Supplementary Figure S5**

Characterization of cytokine fusions. **A,B**, SDS-PAGE of (**A**) Nickel-NTA purified MSA cytokine fusions and (**B**) Protein A purified Fc cytokine fusions. **C**, HEK Blue IL-12 cells were used to show maintenance of IL-12 signaling capacity of novel cytokine fusions. **D**, CTLL-2 cells were exposed to varying concentrations of IL-15 fusion constructs and their proliferation assessed 48 hours later with Cell Titer Glo 2.0. **E**, Collagen I ELISA of lumican fusions show retention of binding in new formats. **F-I**, Biolayer interferometry of (**F**) IL12-versican S139G$_T$-MSA, (**G**) versican S139$G_T$-MSA-IL15, (**H**) IL12-versican S139$G_T$-Fc, and (**I**) versican S139$G_T$-Fc-IL15 demonstrating maintenance of hyaluronic acid binding.

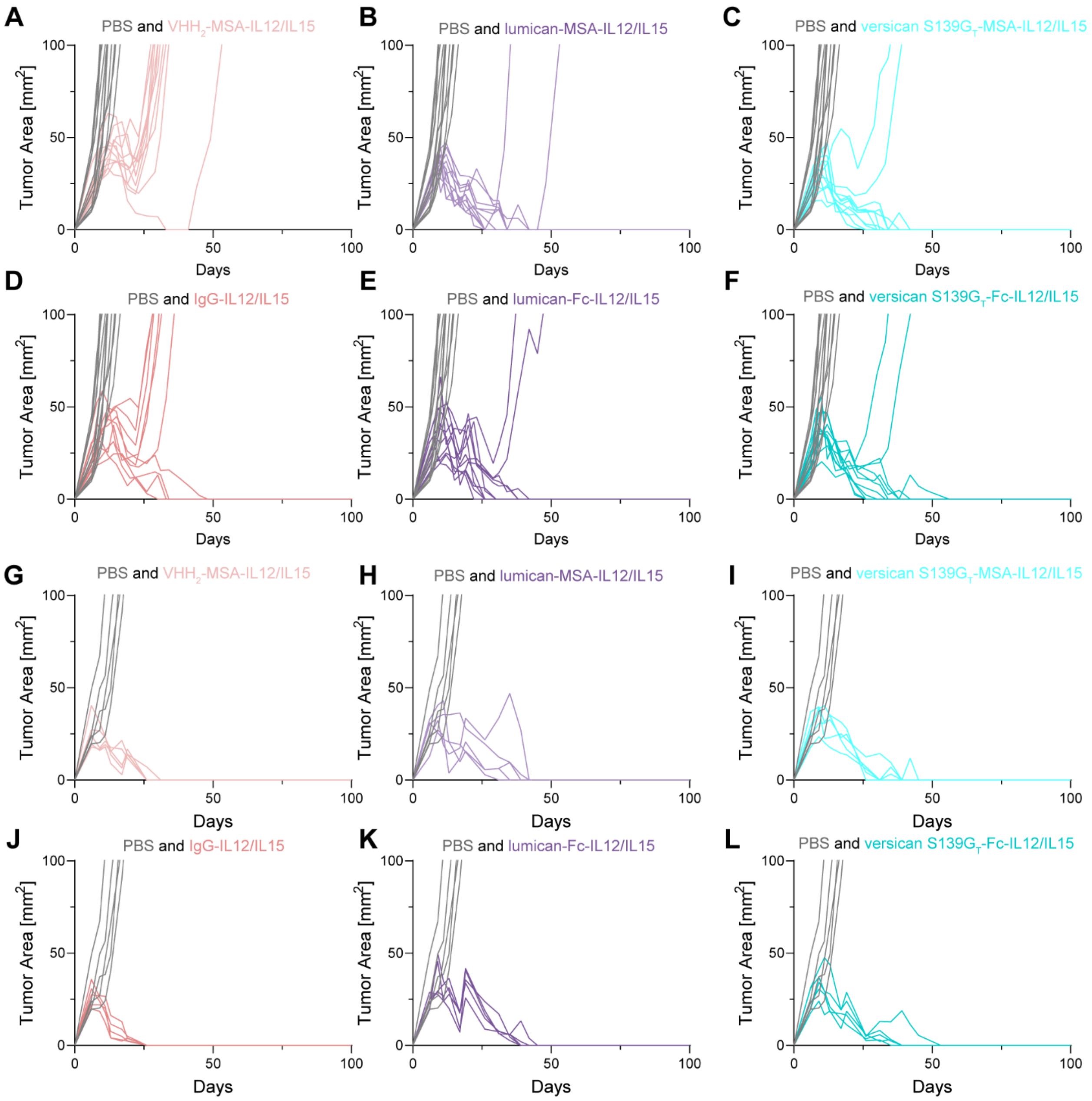


**Supplementary Figure S6**

HA- and collagen-anchored IL-12/IL-15 combination therapy induce regression of B16F10 and MC38 tumors. **A-F**, B16F10 tumor-bearing mice were treated on days 6 and 13 with indicated therapies. B16F10 tumor growth plots of mice receiving PBS, (**A**) $VHH_2$-MSA-IL12/IL15, (**B**) lumican-MSA-IL12/IL15, (**C**) versican $S139G_T$-MSA-IL12/IL15, (**D**) IgG-IL12/IL15, (**E**) lumican-Fc-IL12/IL15, and (**F**) versican $S139G_T$-Fc-IL12/IL15. **G-L**, MC38-tumor bearing mice were treated on days 6 and 13 with the indicated therapies. Tumor growth plots of mice receiving PBS, (**G**) $VHH_2$-MSA-IL12/IL15, (**H**) lumican-MSA-IL12/IL15, (**I**) versican $S139G_T$-MSA-IL12/IL15, (**J**) IgG-IL12/IL15, (**K**) lumican-Fc-IL12/IL15, and (**L**) versican $S139G_T$-Fc-IL12/IL15.

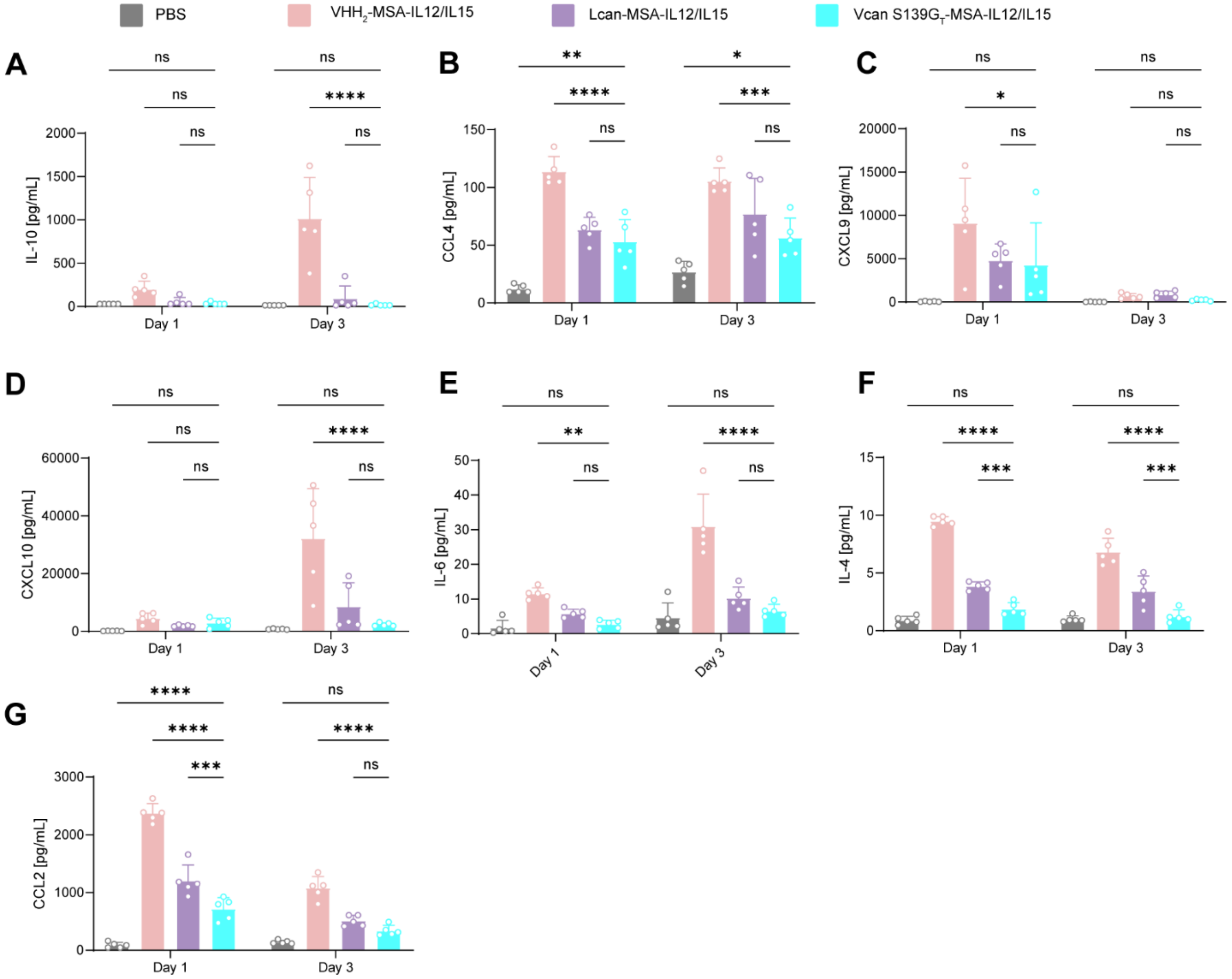


**Supplementary Figure S7**

LegendPlex cytokine release syndrome panel results**. A-G**, Serum concentrations of (**A**) IL-10, (**B**) CCL4, (**C**) CXCL9, (**D**) CXCL10, (**E**) IL-6, (**F**) IL-4, and (**G**) CCL2 as determined by the LegendPlex Mouse Cytokine Release Syndrome panel. Statistics: two-way ANOVA followed by Tukey's multiple-comparison test. ns, not significant; *P < 0.05; **P < 0.01; ***P < 0.001; ****P < 0.0001.

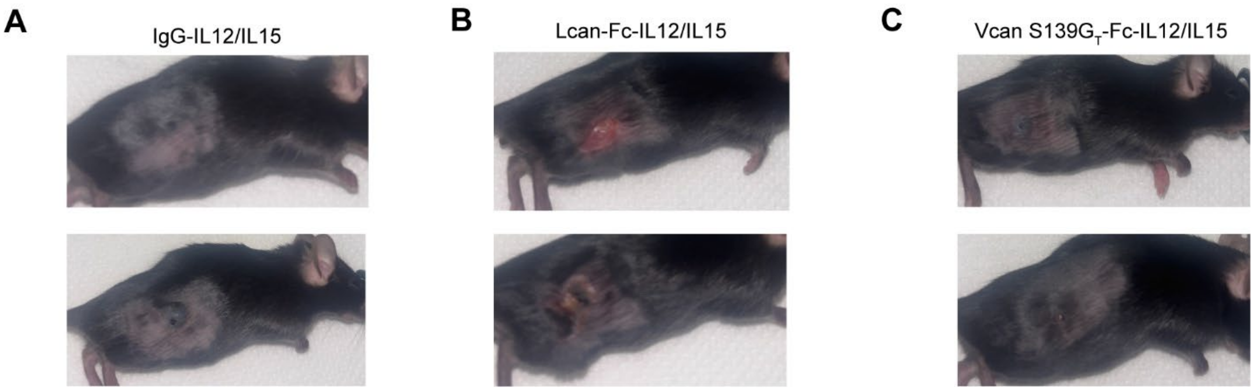


**Supplementary Figure S8**

Mice receiving collagen-anchored cytokine therapy develop skin wounds, where other treatment groups do not. **A-C**, B16F10 tumor-bearing mice were treated with indicated therapies on days 6 and 13 post-tumor induction. Images of the treated area were taken 7 days later. Images of treated area of mice receiving (**A**) IgG-IL12/IL15, (**B**) lumican-Fc-IL12/IL15, or (**C**) versican S139$G_T$-Fc-IL12/IL15 therapy.
